# Interruption of Continuous Opioid Exposure Exacerbates Drug-Evoked Adaptations in the Mesolimbic Dopamine System

**DOI:** 10.1101/646356

**Authors:** Emilia M. Lefevre, Marc T. Pisansky, Carlee Toddes, Federico Baruffaldi, Marco Pravetoni, Lin Tian, Thomas J. Y. Kono, Patrick E. Rothwell

## Abstract

Drug-evoked adaptations in the mesolimbic dopamine system are postulated to drive opioid abuse and addiction. These adaptations vary in magnitude and direction following different patterns of opioid exposure, but few studies have systematically manipulated the pattern of opioid administration while measuring neurobiological and behavioral impact. We exposed male and female mice to morphine for one week, with administration patterns that were either intermittent (daily injections) or continuous (osmotic minipump infusion). We then interrupted continuous morphine exposure with either naloxone-precipitated or spontaneous withdrawal. Continuous morphine exposure caused tolerance to the psychomotor-activating effects of morphine, whereas both intermittent and interrupted morphine exposure caused long-lasting psychomotor sensitization. Given links between locomotor sensitization and mesolimbic dopamine signaling, we used fiber photometry and a genetically encoded dopamine sensor to conduct longitudinal measurements of dopamine dynamics in the nucleus accumbens. Locomotor sensitization caused by interrupted morphine exposure was accompanied by enhanced dopamine signaling in the nucleus accumbens. To further assess downstream consequences on striatal gene expression, we used next-generation RNA sequencing to perform genome-wide transcriptional profiling in the nucleus accumbens and dorsal striatum. The interruption of continuous morphine exposure exacerbated drug-evoked transcriptional changes in both nucleus accumbens and dorsal striatum, dramatically increasing differential gene expression and engaging unique signaling pathways. Our study indicates that opioid-evoked adaptations in brain function and behavior are critically dependent on the pattern of drug administration, and exacerbated by interruption of continuous exposure. Maintaining continuity of chronic opioid administration may therefore represent a strategy to minimize iatrogenic effects on brain reward circuits.

## INTRODUCTION

Opioid analgesics taken for pain relief also activate opioid receptors in the mesolimbic dopamine system, including the ventral tegmental area (VTA) and nucleus accumbens [1,2]. Stimulation of these receptors is positively reinforcing [3,4] and enhances mesolimbic dopamine release in rodents [5,6], though this latter effect has been difficult to detect in humans [7,8]. While dopamine manipulations have mixed effects on acute opioid reward [6,9], chronic opioid exposure produces transcriptional and epigenetic changes in the nucleus accumbens, leading to structural and functional circuit remodeling that are hypothesized to promote addiction and vulnerability to relapse [10–12]. Prevention strategies that minimize these iatrogenic effects might facilitate safer opioid use for clinical indications [13].

Different patterns of drug exposure produce diverse effects on mesolimbic dopamine function and addiction-related behavior (e.g., [14–25]). Decreased sensitivity (i.e., tolerance) is typically reported after relatively continuous patterns of opioid administration, whereas increased sensitivity (i.e., sensitization) is commonly observed after more intermittent patterns of opioid exposure [26–28]. These divergent effects of continuous and intermittent opioid exposure have been reported for mesolimbic dopamine release [29–31], drug reward [32–38], and psychomotor activation [39–43]. Given the variable pharmacokinetics of prescription opioid formulations purported to provide continuous action [44], it is critical to understand how the pattern of opioid exposure shapes adaptations in the mesolimbic dopamine system, in order to minimize/prevent adaptations that promote opioid abuse and addiction. Based on prior literature [45–48], our guiding hypothesis was that maintaining the continuity of opioid exposure reduces iatrogenic effects, by preventing withdrawal caused by fluctuating drug levels.

To test this hypothesis, we first replicated prior reports that intermittent morphine injections produce psychomotor sensitization, whereas continuous morphine infusion produces psychomotor tolerance. However, this comparison was confounded by large differences in pharmacokinetic variables like cumulative dose and peak drug level. We therefore adopted a pharmacodynamic strategy of interrupting continuous morphine administration with daily injections of an opioid receptor antagonist, to precipitate a state of withdrawal. This manipulation provided control over pharmacokinetic variables, and caused a reversal of psychomotor adaptation from tolerance to sensitization. This switch to locomotor sensitization was accompanied by enhanced dopamine signaling in the nucleus accumbens, as well as changes in striatal gene expression measured with next-generation RNA sequencing. Together, our data suggest sensitization of the mesolimbic dopamine system can be minimized by maintaining the continuity of opioid exposure during chronic treatment, highlighting an actionable prevention strategy to reduce opioid abuse.

## MATERIALS AND METHODS

### Subjects

Male and female C57BL/6J mice, mu opioid receptor (*Oprm1*) knockout mice [49], and dopamine transporter (DAT)-IRES-Cre knock-in mice [50] were obtained from The Jackson Laboratory or bred in-house. Experimental procedures were approved by the Institutional Animal Care and Use Committee of the University of Minnesota. For additional details, see Supplementary Information.

### Drug Exposure

Morphine hydrochloride (Mallinckrodt) was dissolved in sterile saline (0.9%), and delivered subcutaneously by bolus injection (5 mL/kg), continuous infusion using osmotic minipumps (Alzet Model 2001), or programmed infusion using miniaturized mechanical pumps (iPrecio SMP-300). For additional details, see Supplementary Information. To interrupt continuous morphine exposure, we injected mice with saline or naloxone (0.1-10 mg/kg, s.c.) twice per day, with injections separated by a period of two hours [51]. Behavioral assessments were performed prior to naloxone injection on the first day of morphine exposure, and again 24 hours after the final naloxone injection.

### Behavioral and Pharmacokinetic Assessments

We tested open-field locomotor activity in a clear plexiglass arena (ENV-510, Med Associates) housed within a sound-attenuating chamber. Thermal antinociception was tested on a 55°C hot plate (IITC Life Scientific). Serum morphine were measured by gas chromatography coupled with mass spectrometry as previously described [52,53]. For additional details, see Supplementary Information.

### Stereotaxic Surgery and Fiber Photometry

Intracranial virus injection and optic fiber implantation were performed as previously described [54]. Continuous fiber photometry recordings were conducted in the open-field chambers described above, for 30 minutes before and after naloxone injection on Days 1 and 7, with spontaneous fluorescent transient events detected as previously described [55]. On challenge days, recordings were conducted for 30 minutes before and after injection of morphine or fentanyl. The average fluorescent signal following challenge injection was compared with baseline prior to injection. For additional details, see Supplementary Information.

### Gene Expression and RNA Sequencing

RNA sequencing was performed using male mice to minimize variability, while equal numbers of both sexes were used in all other experiments, including targeted gene expression analysis with qPCR. Following six days of chronic treatment, we rapidly removed brains under isoflurane anesthesia, and dissected bilateral nucleus accumbens (core and shell) and dorsal striatum (caudate-putamen) on ice. For additional details, see Supplementary Information.

### Statistical Analyses

Analysis of variance (ANOVA) was conducted in IBM SPSS Statistics v24, with details provided in Supplemental Information. In the text, we report significant effects that are critical for data interpretation, but comprehensive reporting of all main effects and interactions from ANOVA models can be found in Table S2. Significant simple effects within group are indicated by a hash (#) to the right of group data, while significant simple effects or post-hoc tests between groups are indicated by an asterisk (*) above the data. All summary data are displayed as mean±SEM, with individual data points from male and female mice shown as closed and open symbols, respectively.

## RESULTS

### Behavioral Effects of Intermittent Injection versus Continuous Infusion of Morphine

To compare different patterns of opioid exposure, we first delivered morphine for one week by daily injection (Figure 1A) or continuous infusion (Figure 1E), and measured open field locomotor activity on the first and last day. Morphine caused a dose-dependent increase in locomotion after both injection (Figure 1B; main effect of Dose: F_4,62_=94.91, p<0.001) and infusion (Figure 1F; main effect of Dose: F_2,42_=40.83, p<0.001). Daily injections caused psychomotor sensitization (Figure 1C; Dose × Day interaction F_4,62_=46.84, p<0.001), whereas continuous infusion caused psychomotor tolerance, (Figure 1G; Dose × Day interaction: F_2,42_=8.94, p=0.001), while both exposure patterns caused antinociceptive tolerance (Figure S1A-B). To examine the persistence of psychomotor adaptation, we challenged all groups with morphine injections 10 days after the end of chronic treatment. Psychomotor sensitization persisted during this challenge test (Figure 1D; Pretreatment Dose × Challenge Dose interaction: F_9.92,153.75_=8.55, p<0.001), whereas psychomotor tolerance did not persist (Figure 1H; Pretreatment Dose × Challenge Dose interaction: F_5.75,51.73_=2.00, p=0.085). The acute response to 6.32 mg/kg morphine (Figure 1B) was greater than the challenge response to this dose, likely reflecting novelty of the testing environment on Day 1.

**Figure 1.**
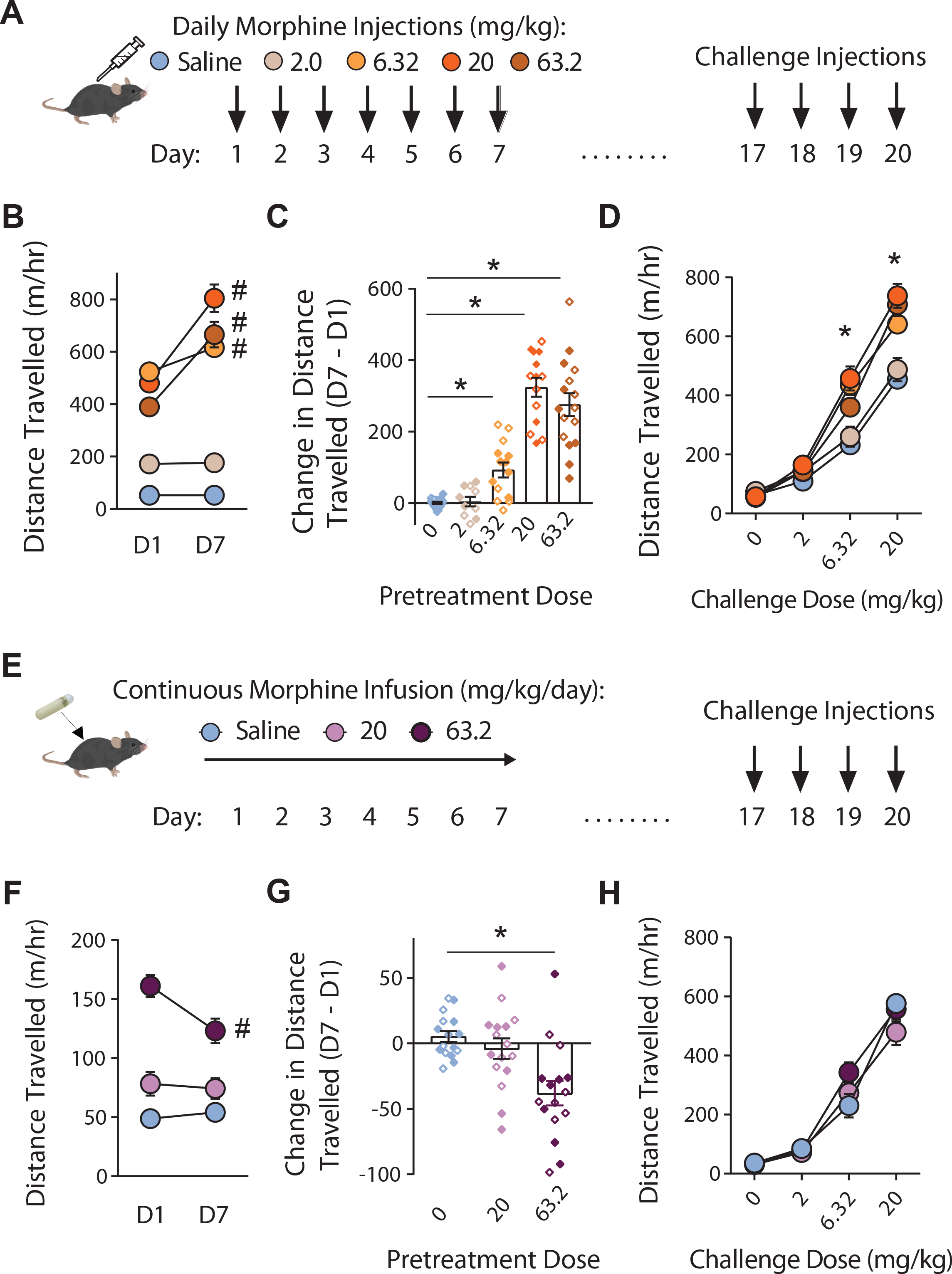
Behavioral effects of intermittent injection versus continuous infusion of morphine. **(A)** Intermittent exposure consisted of seven daily injections of morphine, followed 10 days later by challenge with injection of escalating morphine doses (n=10-18/group). **(B)** Locomotor activity after the first (D1) and last (D7) day of exposure. **(C)** Change in locomotor activity on D7 versus D1, depicted for individual mice at each dose. **(D)** Locomotor activity following challenge injections with ascending doses of morphine. **(E)** Continuous infusion of morphine via osmotic minipump for seven days, followed 10 days later by challenge with injection of escalating morphine doses (n=8/group). **(F)** Locomotor activity on the first (D1) and last (D7) day of exposure. **(G)** Change in locomotor activity on D7 versus D1, depicted for individual mice at each dose. **(H)** Locomotor activity following challenge injections with ascending doses of morphine. All groups contained similar numbers of female mice (open symbols) and male mice (closed symbols); see Supplementary Table 2 for detailed statistical analyses. *p<0.05 between groups, LSD post-hoc test. ^#^p<0.05 for simple effect within group.

These results show that different patterns of chronic morphine exposure cause similar adaptations in some behavioral responses (e.g., antinociception), but divergent adaptations in other behaviors (e.g., psychomotor activation), even when controlling for cumulative dose (e.g., 63.2 mg/kg/day). However, serum morphine levels were predictably higher after bolus injection versus continuous infusion (Figure S1C-D), confounding the effect of exposure pattern with differences in the location/proportion of activated opioid receptors. This highlights the difficulty in simultaneously controlling multiple pharmacokinetic variables (e.g., cumulative dose and peak drug level) when comparing injections and infusions, motivating us to develop a model providing better control of these variables.

### Behavioral Effects of Continuous versus Interrupted Morphine Exposure

To interrupt continuous morphine infusion while controlling pharmacokinetic variables, we administered two daily naloxone injections separated by an interval of two hours (Figure 2A), as previously described [51]. We selected a high dose of naloxone (10 mg/kg) to fully interrupt activation of opioid receptors by morphine, as pilot studies showed this naloxone dose had more robust effects than lower doses (Figure S2). The state of withdrawal precipitated by this dose of naloxone did not significantly change in severity across days (Figure S3).

**Figure 2.**
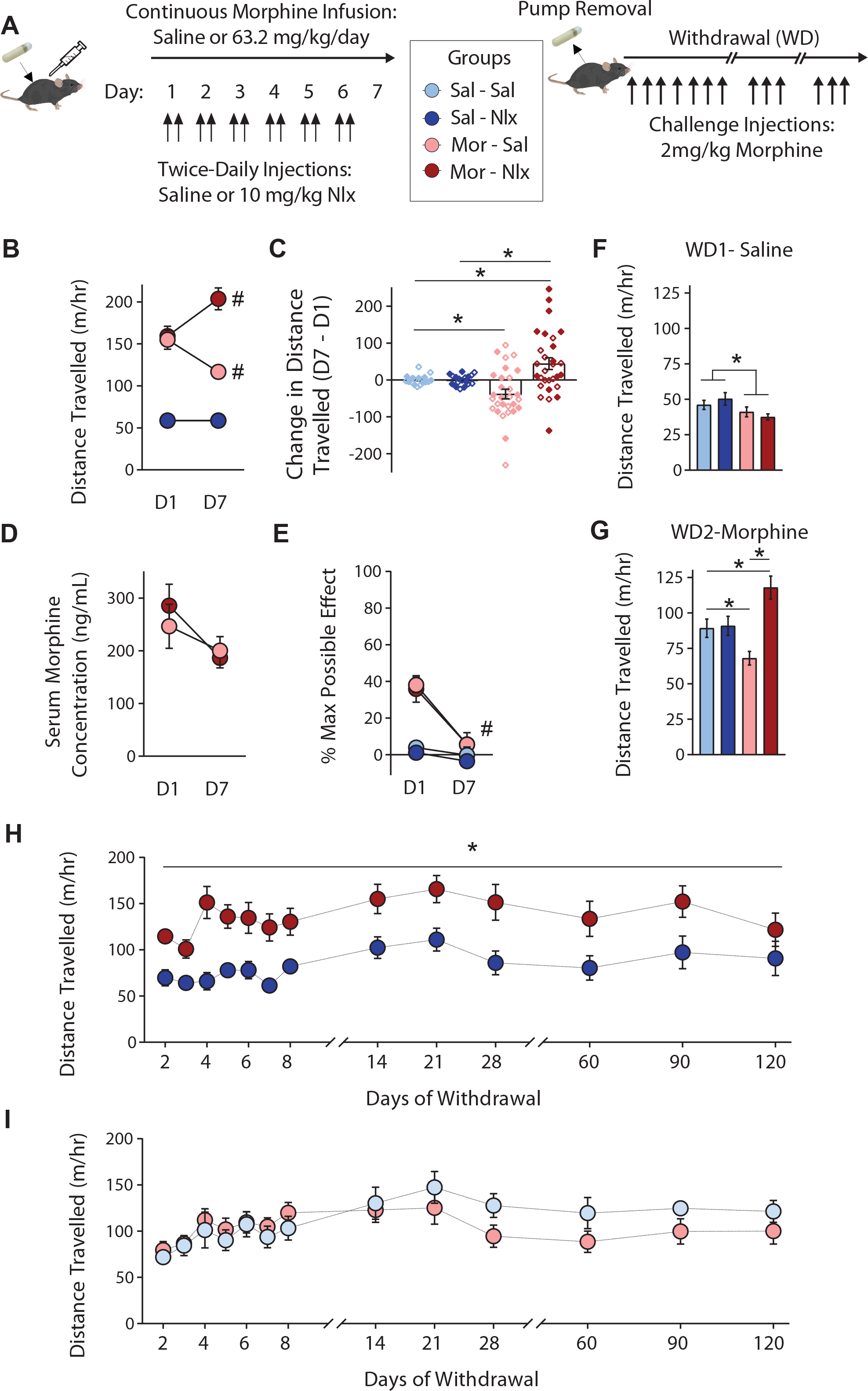
Interruption continuous morphine exposure with daily naloxone injections. **(A)** Continuous infusion of morphine or saline for one week, interrupted by twice-daily injections (separated by 2 hours) of saline or naloxone, followed by challenge injections of morphine (n=18-29/group). **(B)** Locomotor activity on the first (D1) and last (D7) day of exposure; note that data points from Sal-Sal and Sal-Nlx groups are overlaid. **(C)** Change in locomotor activity on D7 versus D1, depicted for individual mice at each dose. *p<0.05 between groups, Tukey post-hoc test. **(D)** Serum morphine concentrations taken on D1 and D7 (n=5/group). **(E)** Thermal antinociception on the hot plate, measured on D1 and D7. ^#^p<0.05 for simple effect within group. **(F)** Open field locomotor activity 24 hours following pump removal. *p<0.05, main effect of Morphine. **(G)** Locomotor response to morphine challenge 48 hours after pump removal. *p<0.05 between groups, LSD post-hoc test. **(H-I)** Locomotor response to morphine challenges during extended withdrawal in groups that previously received daily injections of naloxone (**H**) or saline (**I**) (n=12/group). *p<0.05, main effect of Morphine. All groups contained similar numbers of female mice (open symbols) and male mice (closed symbols); see Supplementary Table 2 for detailed statistical analyses.

In open field tests of locomotor activity (Figure 2B), control mice receiving continuous morphine exposure with saline injections displayed psychomotor tolerance. In contrast, mice receiving naloxone injections to interrupt morphine exposure displayed a switch to psychomotor sensitization (Morphine × Naloxone × Day interaction: F_1,86_=10.32, p=0.002). The increase in locomotion on Day 7 versus Day 1 (Figure 2C) was numerically larger in males (71.64+29.11 m/hr) than in females (17.12+12.25 m/hr), although there were no statistically significant sex differences (Table S2). Serum morphine levels were equivalent between groups on the last day of chronic morphine exposure (Figure 2D), and the development of antinociceptive tolerance was not affected by naloxone injections (Figure 2E). Interrupted morphine exposure had no behavioral effect in mu opioid receptor knockout mice (Figure S4), as previously reported for intermittent morphine injections [49,56]. Psychomotor tolerance was also reversed when continuous morphine administration was interrupted by periods of spontaneous withdrawal, using miniaturized programmable infusion pumps (Figure S5).

Twenty-four hours after pump removal, locomotion was decreased in both morphine groups, reflecting a state of spontaneous withdrawal (Figure 2F; main effect of Morphine: F_1,40_=12.16, p=0.001). The next day, all mice were challenged with injection of 2 mg/kg morphine (Figure 2G). Psychomotor activation was blunted by previous exposure to continuous morphine, while sensitization persisted after interrupted morphine (Morphine × Naloxone interaction: F_1,64_=13.37, p=0.001), suggesting both effects last for 48 hours after chronic exposure. To map the persistence of these effects, a subset of mice continued receiving 2 mg/kg morphine challenges. We again observed a significant Morphine × Naloxone interaction (F_1,39_=13.97, p<0.001), and thus separately analyzed the simple effect of morphine in each injection group. After interrupted morphine, psychomotor sensitization was evident in response to daily and weekly challenges, but then gradually diminished over subsequent months (Figure 2H; main effect of Morphine: F_1,17_=14.21, p=0.002). After continuous morphine, psychomotor tolerance dissipated almost immediately (Figure 2I; main effect of Morphine: F_1,18_=1.23, p=0.28).

### Mesolimbic Dopamine Signaling after Continuous versus Interrupted Morphine Exposure

To determine whether these divergent behavioral changes are associated with different patterns of dopamine signaling in the nucleus accumbens, we used fiber photometry to measure fluorescent signals from a genetically encoded dopamine sensor, dLight1.3b [57,58]. We stereotaxically injected adeno-associated virus (AAV) expressing dLight1.3b into the nucleus accumbens, followed by optic fiber implantation above the site of virus injection (Figure 3A). This experiment included some DAT-IRES-Cre knock-in mice, which received a second VTA injection of AAV expressing Cre-dependent ChrimsonR, a red-shifted excitatory opsin [59] (Figure 3A). ChrimsonR was expressed by dopamine neurons in the VTA and transported to their axon terminals in nucleus accumbens (Figure 3B). We delivered red light (595nm) to stimulate dopamine release, and monitored dLight1.3b signal through the same optic fiber at both dopamine-dependent and control wavelengths (470/405nm, respectively; Figure 3C). To validate the correspondence between dLight1.3b fluorescence and dopamine signaling, we confirmed that the fluorescent signal increased with red-shifted optogenetic stimulation in a frequency-dependent fashion (Figure 3D), and was blocked by the D1 dopamine receptor antagonist SCH23390 (Figure 3E).

**Figure 3.**
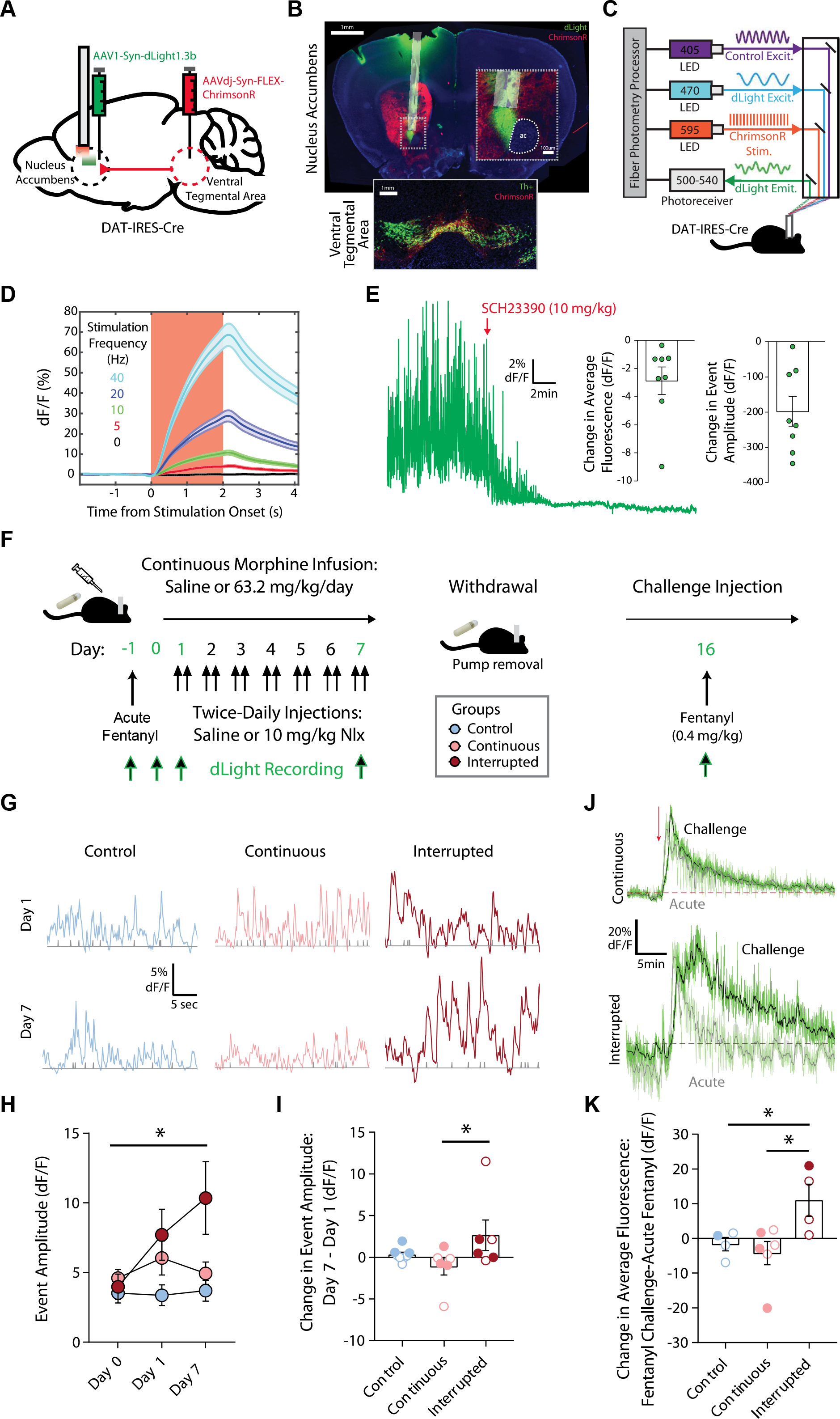
Mesolimbic dopamine sensitization induced by interrupted morphine. **(A)** Stereotaxic injection of AAV1-Syn-dLight1.3b into the nucleus accumbens and AAVdj-Syn-FLEX-ChrimsonR into the VTA of DAT-IRES-Cre mice (n=10). **(B)** Top panel: Example of optic fiber implanted in the nucleus accumbens core, with local dLight expression (green) and ChrimsonR expression in dopamine axon terminals (red). Bottom panel: ChrimsonR expression in the VTA, with dopamine neurons labelled by tyrosine hydroxylase (Th) immunofluorescence. Scale bar = 1mm. **(C)** Setup for simultaneous optogenetic stimulation (595 nm) and fiber photometry recording (405/470 nm). **(D)** Frequency-dependent changes in dLight fluorescent signal following optogenetic stimulation of ChrimsonR dopamine terminals in the nucleus accumbens. **(E)** Trace showing block of fluorescent signal by the D1 receptor antagonist SCH23390 (red arrow denotes s.c. injection). Inset: quantification of SCH23390 effects on average fluorescent signal (left) and transient event amplitude (right). (F) Experimental timeline for photometry recordings. **(G)** Examples of spontaneous fluorescent transients recorded on days 1 and 7. **(H)** Average transient event amplitude on days 0, 1 and 7 for Control, Continuous and Interrupted morphine treatment groups (n=6-8/group). *p<0.05, Group × Day interaction. **(I)** Change in transient event size on D7 versus D1, depicted for individual mice. *p<0.05, LSD post-hoc test. **(J)** Representative trace for showing the response to acute fentanyl (light green) and fentanyl challenge (dark green), for continuous morphine (top) and interrupted morphine (bottom). **(K)** Change in average fluorescent signal after fentanyl challenge, compared to acute fentanyl. *p<0.05, LSD post-hoc test; note that challenge data from six mice are missing due to lost head caps. All groups contained similar numbers of female mice (open symbols) and male mice (closed symbols); see Supplementary Table 2 for detailed statistical analyses.

We next conducted fiber photometry recordings the day prior to implantation of osmotic pumps, and on Days 1 and 7 of chronic morphine treatment (Figure 3F). Naloxone injection significantly reduced fluorescent signals on both Days 1 and 7 in mice implanted with morphine pumps, but not mice implanted with saline pumps (Figure S3C-F). There were no differences in fluorescent signals between mice implanted with saline pumps and injected with either saline or naloxone, so these groups were combined to form a single “control” group. To quantify changes in baseline dopamine dynamics, we used peak analysis to identify spontaneous fluorescent transient events [60], which were blocked by SCH23390 (Figure 3E). The amplitude of these transient events was increased in both morphine groups prior to naloxone injection on Day 1, but then diverged on Day 7 (Figure 3H; Group × Day interaction; F_4,28_=5.20, p=0.006). Event amplitude tended to decrease over the course of continuous morphine exposure, but tended to increase over the course of interrupted morphine exposure (Figure 3I; main effect of Group: F_2,14_=2.71, p=0.10).

To examine the persistence of these changes in dopamine signaling, we administered a challenge injection of morphine (2 mg/kg) one week after pump removal, a time point where interrupted morphine treatment caused locomotor sensitization (Figure 2H). The interrupted morphine group showed an increase in average fluorescent signal after morphine challenge, which tended to be larger than either the control or continuous morphine group (Figure S6). However, the change in fluorescent signal was modest and gradual, most likely because of morphine’s slow pharmacokinetics relative to other opioids. These mice received an acute injection of fentanyl (0.4 mg/kg) prior to chronic morphine treatment (Day −1), which generated a rapid and robust signal similar to heroin [6], so we challenged them with the same dose of fentanyl one day after morphine challenge (Figure 3J). The interrupted morphine group showed a significant enhancement in the response to fentanyl challenge, compared to acute fentanyl injection (Figure 3K; main effect of Group: F_2,8_=6.23, p=0.023). Overall, these data provide convergent evidence for persistent sensitization of the mesolimbic dopamine system following interrupted morphine exposure.

### Differential Gene Expression after Continuous versus Interrupted Morphine Exposure

We next assessed the downstream consequences of continuous and interrupted morphine exposure on gene expression in the nucleus accumbens. To minimize variability related to sex differences, we used only male mice in this experiment, since interrupted morphine caused more robust locomotor sensitization in males. At the end of chronic treatment, we dissected the nucleus accumbens as well as the dorsal striatum (Figure 4A), and used next-generation RNA sequencing to perform genome-wide transcriptional profiling. We defined differential gene expression with a fold change threshold of 15%, while controlling false discovery rate at q<0.05. With these criteria, there was no evidence of differential gene expression between saline-saline and saline-naloxone treatments (Table S3), so these treatments were combined to form a single “control” group.

**Figure 4.**
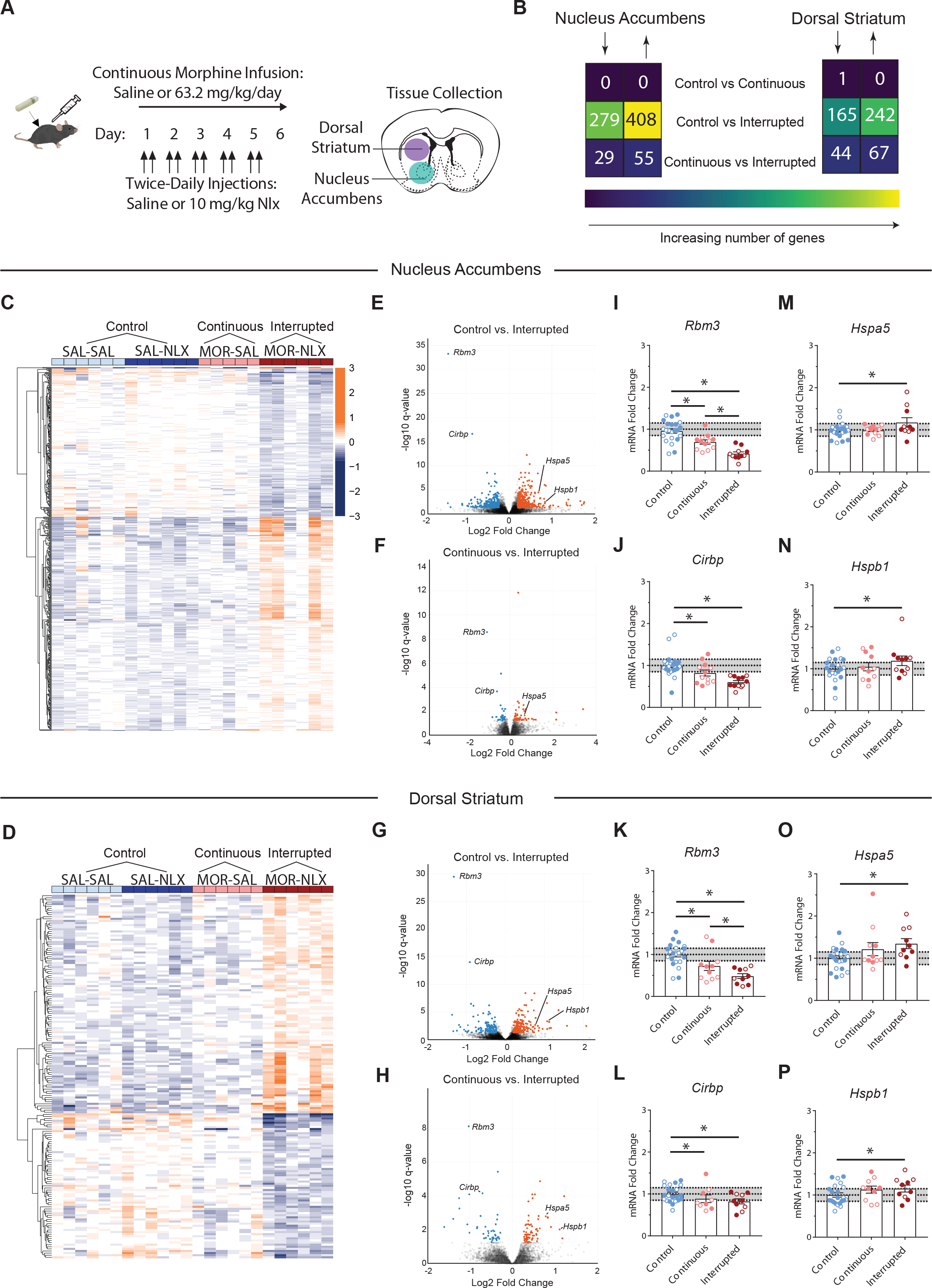
Differential gene expression after continuous or interrupted morphine. **(A)** Microdissection of nucleus accumbens or dorsal striatum tissue for RNA sequencing after six days of continuous or interrupted morphine exposure (n=5-6 male mice/group). **(B)** The number of differentially expressed genes (DEGs) that were significantly up- or down-regulated in each brain region; note that mice implanted with saline pumps have been combined to form a single control group. **(C-D)** Heat maps showing normalized level of DEGs for each individual sample in the nucleus accumbens (C) and dorsal striatum (D). **(E-H)** Volcano plots of significantly
up- and down-regulated genes after interrupted morphine in the nucleus accumbens (E-F) and dorsal striatum (G-H). Highlighted are cold-shock proteins Rbm3 (RNA binding motif protein 3) and Cirbp (cold inducible RNA binding protein) and heat-shock proteins Hspb1 and Hspa5. **(I-P)** Independent validation of highlighted genes with quantitative RT-PCR. Shaded area shows the 15% fold change threshold used to define differential gene expression in RNA sequencing data (n=10-24/group), with similar numbers of female mice (open symbols) and male mice (closed symbols) for PCR validation; see Supplemental Table S2 for detailed statistical analyses. *p=0.05, LSD post-hoc test.

Compared to the control group, interrupted morphine significantly regulated 687 transcripts in the nucleus accumbens and 407 transcripts in the dorsal striatum (Figure 4B-D). Surprisingly, continuous morphine significantly regulated only one gene (*Sst*) in the dorsal striatum. With a less stringent statistical threshold of p<0.05, continuous morphine significantly regulated 112 genes in the nucleus accumbens and 294 genes in the dorsal striatum (Figure S7), comparable to previous studies of chronic morphine exposure [61,62]. Interrupted morphine still had substantially greater impact, significantly regulating 1,389 and 1,382 genes in the nucleus accumbens and dorsal striatum, respectively. Using our original statistical criteria (q<0.05), direct comparison of continuous and interrupted morphine (Figure 4B) also revealed differential gene expression in both the nucleus accumbens (84 transcripts) and dorsal striatum (111 transcripts).

In both the nucleus accumbens and dorsal striatum (Figure 4E-H), there was a high degree of similarity in the effect of interrupted morphine compared to either continuous morphine or control (Figure S8). There were no significant changes in expression of opioid or dopamine receptors, but a number of downstream signaling molecules showed significant regulation following interrupted morphine exposure: G-protein subunits (Gna11/Gnaz/Gnb2), regulators of G-protein signaling (Rgs2/Rgs11/Rgs20), adenylate cyclase (Adcy8), phosphodiesterase (Enpp6), and transcription factors (Creb1/Crebl2). Several of these genes have previously been shown to regulate responses to morphine [63–72]. Two of the most dramatic effects were downregulation of Rbm3 and Cirbp, which were previously reported to be downregulated in the nucleus accumbens following continuous morphine exposure [61]. We used quantitative RT-PCR to validate these changes in tissue from an independent cohort of male and female mice. In both nucleus accumbens and dorsal striatum, Rbm3 and Cirbp expression was decreased after continuous morphine and further reduced by interrupted morphine (Figure 4I-L). We also validated the upregulation of two heat shock proteins, Hspb1 and Hspa5, after interrupted morphine exposure (Figure 4M-P).

### Ingenuity Pathway Analysis and Weighted Gene Coexpression Network Analysis

Using differentially expressed gene lists, we performed unbiased identification of signaling pathways regulated by interrupted morphine exposure using Ingenuity Pathway Analysis. Despite significant regulation of individual genes involved in opioid and dopamine receptor signaling, the “Opioid Signaling” and “Dopamine Receptor Signaling” canonical pathways were not significantly enriched (Table S4). Instead, several unexpected canonical pathways were significantly regulated in all comparisons of interest across both striatal subregions (Figure 5A-B). The majority of individual molecules driving significant changes in these pathways were heat shock proteins (Table S4). In comparison to both control and continuous morphine, interrupted morphine upregulated transcripts encoding numerous individual heat shock proteins in the nucleus accumbens and dorsal striatum (Figure S9). Significant changes in two top upstream regulators, heat shock transcription factor 1 (HSF1) and sphingosine-1-phosphate phosphatase 2 (SGPP2), were also driven primarily by heat shock proteins (Table S5).

**Figure 5.**
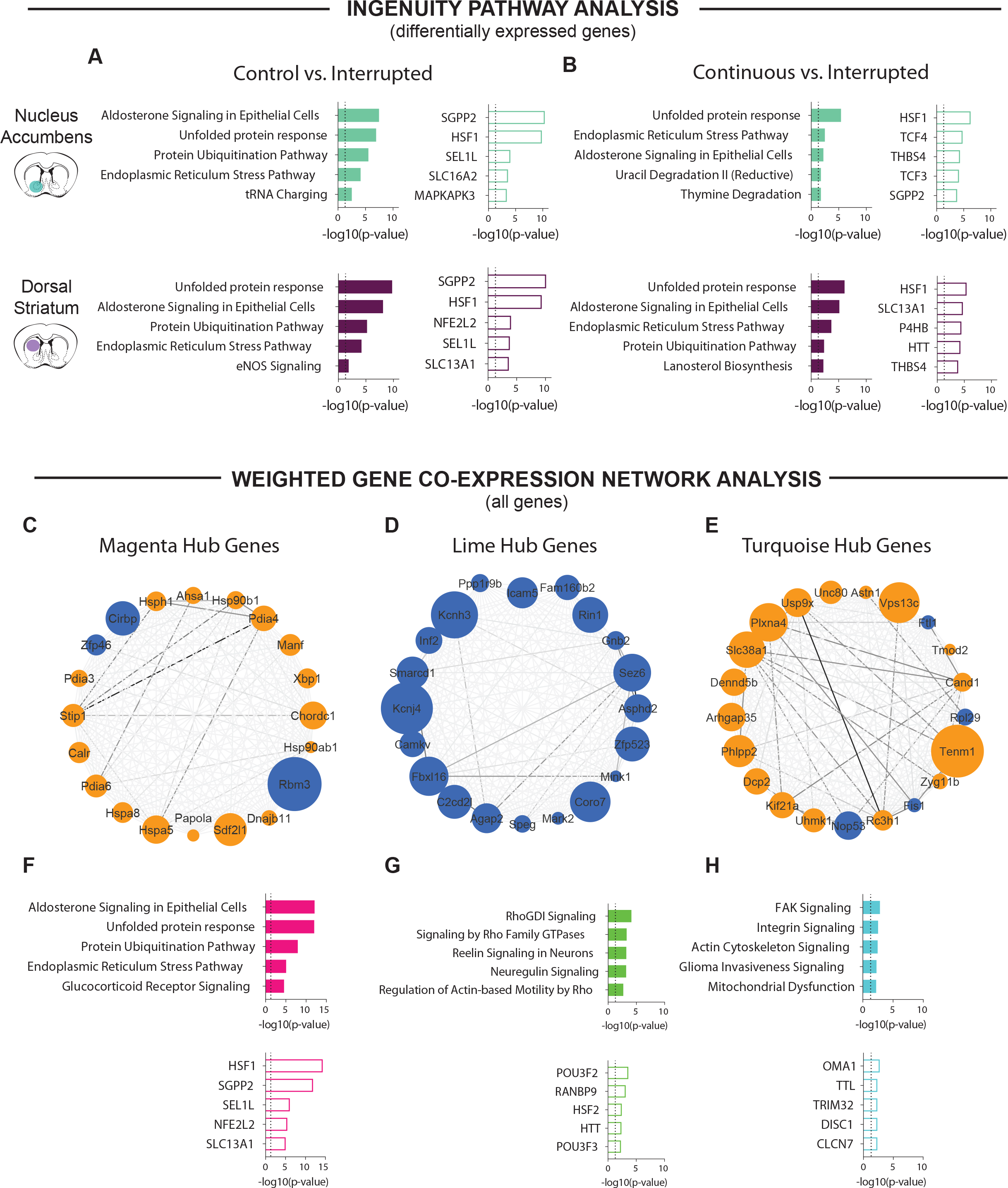
Characterization of differentially expressed genes. **(A-B)** Top five canonical pathways (closed bar) and upstream regulators (open bars) identified by Ingenuity Pathway Analysis for interrupted morphine versus control (A) or continuous morphine (B), in the nucleus accumbens (top row) and dorsal striatum (bottom row). Dotted line on bar graphs indicates p=0.05; for details regarding drug treatment, refer Figure 4 (n=5-12 male mice/group). **(C-E)** Weighted gene co-expression network analysis identified three consensus modules across the nucleus accumbens and dorsal striatum: magenta (C), lime (D) and turquoise (E). Top 20 hub genes with the highest degree of connectivity are displayed for each module, with node size scaled by fold change in the control vs. interrupted morphine comparison, and node color indicating up-regulation (orange) or down-regulation (blue). The weight of edges connecting nodes is scaled such that stronger connections are darker in color. **(F-H)** Ingenuity Pathway Analysis of canonical pathways (top row) and upstream regulators (bottom row) for genes in the magenta module (F), lime module (G), and turquoise module (H).

We further analyzed RNA sequencing data using weighted gene co-expression network analysis (WGCNA), a computational method that does not require binary thresholds for differential gene expression [73]. We identified 18 consensus modules that exhibited correlated patterns of expression across samples and treatment conditions, which were given arbitrary color names (Figure S10A). The magenta module showed significant regulation in both nucleus accumbens and dorsal striatum, while the turquoise and lime modules were only significantly regulated in the nucleus accumbens (Figure S10B-E). We generated connectivity diagrams to visualize “hub” genes with the highest degree of connectivity within each module (Figure 5C-E and Table S6).

Hub genes in the magenta module included Cirbp and Rbm3, and the two hub genes with greatest connectivity were heat shock proteins, Hspa5 and Hspa8 (Figure 5E). Many of the same canonical pathways and upstream regulators detected in our analysis of differentially expressed genes were also represented in the magenta module (Figure 5F). The turquoise and lime modules (Figure 5G-H) implicated additional signaling pathways in the nucleus accumbens, including upregulation of FAK and actin cytoskeleton signaling (turquoise module) [74], as well as downregulation of signaling by Rho family GTPases (lime module). These results provide convergent evidence that interruption of continuous morphine exposure exacerbates drug-evoked changes in striatal gene expression, engaging novel signaling pathways that may contribute to opioid abuse and addiction.

## DISCUSSION

Our data provide clear evidence that the pattern of opioid administration dictates drug-evoked adaptations in the mesolimbic dopamine system. Using psychomotor activity as a behavioral readout [75], we first replicated prior work showing robust psychomotor sensitization after intermittent morphine injections [39–43], while continuous morphine infusion caused psychomotor tolerance. The interruption of continuous opioid receptor stimulation with either spontaneous or naloxone-precipitated withdrawal caused a behavioral reversal of psychomotor tolerance. This behavioral switch was associated with enhancements of both dopamine signaling and gene expression in the nucleus accumbens, highlighting the fundamental influence of exposure pattern on opioid-evoked adaptations in this brain region.

### Behavioral and Neurochemical Sensitization after Interrupted Morphine Exposure

The neurobehavioral impact of chronic opioid exposure has been investigated in rodents using a wide variety of drug administration regimens. Our data confirm that daily morphine injections at a fixed dose cause psychomotor sensitization, whereas psychomotor tolerance is observed following either continuous exposure or multiple daily injections at escalating doses [39–43]. These patterns differ in the degree to which drug effects dissipate between each administration, and thus the cumulative amount of withdrawal that occurs over the course of chronic opioid administration. Periods of withdrawal between intermittent opioid exposure may represent a form of stress [46], leading to adverse consequences that can be avoided through sustained opioid receptor activation [48], including methadone maintenance therapy [45].

We used two strategies to interrupt continuous morphine administration, while simultaneously maintaining control of critical pharmacokinetic variables like cumulative dose and peak drug level. The first was a pharmacodynamic manipulation involving daily administration of naloxone to precipitate a state of withdrawal [51], which also represents a form of stress. The second was a pharmacokinetic manipulation, using miniaturized programmable infusion pumps to periodically shut off drug delivery and produce a state of spontaneous withdrawal. Both manipulations completely reversed the development of psychomotor tolerance normally caused by continuous morphine exposure. The psychomotor sensitization that developed after interrupting morphine exposure with naloxone persisted for weeks to months, mirroring the durable sensitization produced by intermittent morphine injections [76,77], and contrasting with the transient nature of psychomotor tolerance [78,79].

The persistence of psychomotor sensitization reflects long-lasting adaptations in nucleus accumbens dopamine and glutamate signaling [75], which also enhance the incentive properties of drugs and associated cues [80]. To measure dynamic changes in nucleus accumbens dopamine signaling, we used fiber photometry to monitor fluorescent signals from a genetically encoded dopamine sensor, dLight1.3b [57,58]. The amplitude of spontaneous fluorescent transients increased in parallel with the development of psychomotor sensitization during interrupted morphine exposure, similar to changes detected using fast-scan cyclic voltammetry following cocaine exposure [81]. One week after interrupted morphine exposure, we also detected an enhanced dopamine response to challenge injection of fentanyl. These data are consistent with enhanced mesolimbic dopamine release reported after intermittent morphine exposure [29,30], and contrast with reduced dopamine release following continuous morphine exposure [31]. These effects on dopamine release may contribute to the respective sensitization and tolerance of drug reward following intermittent and continuous opioid exposure [32–38], although further research is needed to determine whether interrupted opioid exposure increases subsequent sensitivity to opioid reward.

### Nucleus Accumbens Gene Expression after Interrupted Morphine Exposure

The distinct behavioral effects of continuous and interrupted morphine exposure were mirrored at the level of gene expression in the nucleus accumbens and dorsal striatum. In both brain regions, the number of differentially expressed transcripts was substantially greater after interrupted versus continuous morphine. Continuous morphine regulated 100-200 genes by conventional statistical standards (p<0.05), similar to previous reports [61], but only a single gene reached statistical criteria for differential expression after controlling for false discovery rate (q<0.05). The impact of continuous morphine on striatal gene expression is thus less severe than interrupted morphine, as previously reported for chronic morphine treatment and morphine withdrawal in other brain regions [82]. Naloxone injection did not significantly alter gene expression in control mice implanted with saline pumps, most likely because tissue was collected 24 hours after the last naloxone injection, permitting time for transcriptional recovery after naloxone injection in the absence of morphine.

Individual genes involved in opioid and dopamine receptor signaling were differentially regulated by interrupted morphine exposure, although Ingenuity Pathway Analysis did not reveal significant enrichment of these canonical pathways (which are not specifically tailored to genes with enriched expression in striatal tissue). The changes in nucleus accumbens dopamine signaling we report may begin with alterations in dopamine cell bodies in the VTA [31,36], which subsequently impact striatal gene expression. A number of genes that were differentially expressed after interrupted morphine encode synaptic receptors for glutamate (Gria4, Grin2d, Grin3a), GABA (Gabra2, Gabrb2), and glycine (Glra2, Glra3). This is consistent with well-established effects of chronic drug exposure on synaptic transmission in the nucleus accumbens [10], including opioid-evoked plasticity at excitatory and inhibitory synapses [83–85]. Intermittent morphine injections also increase expression of glutamate receptor subunits in the nucleus accumbens [86]. The upregulation of Gabra2 we observe following interrupted morphine contrasts with the downregulation previously reported in the nucleus accumbens after continuous morphine [85], further highlighting the differential impact of these two patterns of exposure.

Ingenuity Pathway Analysis of either differentially expressed gene lists or the WCGNA magenta module identified similar canonical pathways, including unfolded protein response and endoplasmic reticulum stress. Interrupted morphine exposure upregulated many genes encoding heat shock proteins while robustly reducing expression of two cold shock proteins, Rbm3 and Cirbp. Changes in expression of both heat and cold shock proteins have previously been reported in striatal tissue following opioid treatment [61,87–91], and our quantitative RT-PCR analysis shows that changes in expression of these genes are exacerbated by the interruption of continuous morphine exposure with naloxone. These gene expression changes could be related to opioid-induced temperature fluctuations in the nucleus accumbens [92], or a more general response to cellular stress that is independent of temperature. Striatal expression of heat shock proteins is tied to psychomotor sensitization, withdrawal, and other behavioral responses to opioids [89,93–98], supporting functional relevance of the transcriptional changes we observe. As a top upstream transcriptional regulator, HSF1 is a novel and intriguing therapeutic target for addiction that also plays a role in cancer and neurodegenerative disease [99].

In conclusion, our data show that interruption of continuous opioid administration sensitizes mesolimbic dopamine transmission and exacerbates transcriptional changes in the nucleus accumbens and dorsal striatum, producing an enhanced behavioral sensitivity to opioids that persist for months. Maintaining the continuity of chronic opioid administration may therefore represent a strategy to minimize iatrogenic effects on brain reward circuits [13], preventing sensitization of the mesolimbic dopamine system that could otherwise increase vulnerability to subsequent opioid abuse and addiction.

## Supporting information

Figure 4E interactive

Figure 4F interactive

Figure 4G interactive

Figure 4H interactive

## ACKNOWLEDGEMENTS

We thank Kerry Trotter, David Leipold, and Lauren Bystrom for technical assistance, as well as Brian Trieu, Dieter Brandner, and Drs. Cassandra Retzlaff and Rocio Gomez-Pastor for stimulating discussions. Some of the viral vectors used in this study were generated by the University of Minnesota Viral Vector and Cloning Core. The University of Minnesota MnDRIVE Optogenetics Core provided technical support for fiber photometry experiments. The University of Minnesota Genomics Center conducted RNA sequencing, and the Minnesota Supercomputing Institute (MSI) at the University of Minnesota provided resources that contributed to the research results reported within this paper.

