## Supplemental Information for "Interruption of Continuous Opioid Exposure Exacerbates Drug-Evoked Adaptations in the Mesolimbic Dopamine System"

### SUPPLEMENTARY MATERIALS & METHODS

#### Subjects

Male and female C57Bl/6J mice (stock #000664), mu opioid receptor (*Oprm1*) knockout mice (stock #007559), and DAT-IRES-Cre knock-in mice (stock #006660) were obtained from The Jackson Laboratory or bred in-house. Mice were 6-12 weeks old at the beginning of each experiment and housed in groups of 2-5 per cage, on a 12 hour light cycle (0600h – 1800h) at ~23°C with food and water provided ad libitum. Experimental procedures were conducted between 1000h – 1600h.

#### Drug Exposure

Osmotic minipumps (Alzet Model 2001) were implanted in mice weighing up to 25 g, which is the upper limit for administration of 63.2 mg/kg/day morphine using these minipumps. After adjusting morphine concentration for body weight, minipumps were filled with 300 µL of solution and primed overnight at 40°C. Miniaturized programmable infusion pumps (iPrecio SMP-300) were implanted in mice weighing at least 20 g. The pump reservoir was filled with saline or morphine (~50 mg/mL) according to manufacturer's instructions, and then wirelessly programmed to infuse with one of two patterns: (1) a continuous pattern with sustained infusion for 7 consecutive days, or (2) a "discontinuous" pattern with alternating 24-hour periods of drug infusion and pump inactivity for 13 days (see Figure S5). Based on the body weight of each individual mouse, pump management software (iPrecio IMS-300) automatically calculated infusion rates to achieve dosing at 63.2 mg/kg/day for both infusion patterns. The pump reservoir was refilled percutaneously as needed. During the "discontinuous" pattern of morphine administration, mice were weighed daily to confirm weight loss on days the pump was inactive, and two mice failed to show weight loss or rebound on the final two days of the experiment. This likely indicates either a pump malfunction or premature emptying of the pump reservoir, so data from these two mice were excluded from analysis.

All pumps were implanted under anesthesia (5% isoflurane/95% oxygen) through a small incision on the rump, which was then closed with wound clips. Carprofen (5 mg/kg, s.c.) was given as an analgesic before surgery and for 3 days following pump implantation or removal. Behavioral testing began 24 hours after pump implantation (Day 1), to allow for recovery from surgical anesthesia.

**Behavioral and Pharmacokinetic Assessments**

We tested open-field locomotor activity in a clear plexiglass arena (ENV-510, Med Associates) housed within a sound-attenuating chamber. The location of the mouse within the arena was tracked in two dimensions by arrays of infrared beams, connected to a computer running Activity Monitor software (Med Associates). Mice were habituated to the chamber for one hour the day before initiating drug treatment, and then tested on the first and last days of chronic morphine treatment (30 minutes after morphine injections). For morphine challenges, the session length varied as a function of morphine dose: 60 mins (Saline and 2 mg/kg), 90 mins (6.32 mg/kg), or 120 mins (20 mg/kg)

Thermal antinociception was tested on a 55°C hot plate (IITC Life Scientific). The day before initiating drug treatment, mice were habituated to the instrument for 60 seconds at room temperature. Immediately before the first drug exposure, we established baseline latency to either jump or lift and lick a hind paw at 55°C. Mice were then tested on the first and last days of chronic morphine exposure (30 minutes after morphine injections), with a maximal cutoff of 30 seconds to prevent tissue damage. The percent maximum possible effect was calculated as (test latency – baseline latency) / (30 sec – baseline latency) x 100.

In a subset of mice, we also collected facial vein blood samples on the first and last days of morphine exposure (5 minutes after injections), to determine serum morphine concentration. These samples were collected in heparinized tubes and centrifuged at 7500 RPM for 3 minutes at 14°C. Serum was transferred to a 5 ml vial and stored at -20°C until analysis.

A global withdrawal score [1] was assessed on days 1 and 6 of interrupted morphine exposure, during a 10-minute period after the first naloxone injection on that day. Mice were placed in a tall glass cylinder and filmed for offline analysis. An experimenter blind to treatment counted the total number of jumps, rearing and wet dog shakes. Ptosis, rhinorrhea, lacrimation, salivation, diarrhea and eye twitch were evaluated over 2-minute intervals, with a value of 1 being assigned if present during each interval (maximum score 5). Piloerection and locomotor activity were evaluated together over 2-minute intervals, and assigned a value of 0, 1, or 2 for each interval based on minimal, moderate and maximum response, respectively (maximum score 10). All values were added together for the entire 10 minute period to calculate the global withdrawal score.

**Stereotaxic Surgery**

Mice were anesthetized with a ketamine:xylazine cocktail (100:10 mg/kg), and a small hole was drilled above target coordinates for the nucleus accumbens core (AP +1.35, ML +1.10; DV -4.40) and ventral tegmental area (AP -3.0, ML +0.5; DV -4.5). Using a 33-gauge Hamilton syringe, AAV1-Syn-dLight1.3b was delivered to the nucleus accumbens and AAVdj-Syn-FLEX-ChrimsonR-tdTomato was delivered to the ventral tegmental area. Viral solutions were injected at a concentration of ~$10^{13}$ particles/mL, a volume of 0.75 μL, and a rate of 0.1 μL/min. The syringe tip was left in place at the injection site for 5 minutes, and then slowly retracted over the course of 5 minutes. This was followed by unilateral implantation of a 400 μm fiberoptic cannula (Doric Lenses: MFC_400/430-0.48_6mm_MF1.25_FLT) just above the site of virus injection in the nucleus accumbens (+0.02 mm), affixed to the skull using a dual-cure resin (Patterson Dental, Inc.). After surgery, mice were given 500 μL saline and 5 mg/kg carprofen (s.c.) daily for 3 days, and recovered a minimum of 2 weeks before fiber photometry recordings to allow for sufficient virus expression.

**Fiber Photometry Recording and Analysis**

Real-time optical signals were acquired using a RZ5P fiber photometry workstation (Tucker Davis Technologies) as previously described [2]. LEDs at 470 and 405 nm (ThorLabs) were modulated at distinct carrier frequencies (531 Hz and 211 Hz, respectively), and passed through a fluorescence mini cube (Doric Lenses) coupled into a patch cord (400 μm, 0.48 NA). For experiments involving red-shifted optogenetic stimulation with ChrimsonR, a 595 nm LED (ThorLabs) controlled using a pulse generator (Master-8, A.M.P.I.) was connected to the fluorescence mini cube and coupled to the same patch cord. The distal end of the patch cord was connected to the implanted fiberoptic cannula with a ceramic sleeve. Fluorescence was back-projected through the mini cube and focused onto a photoreceiver (Newport Model 2151). Signals were sampled at 6.1 kHz, demodulated in real-time, and saved for offline analysis.

Raw data from each recording session were processed as previously described [3]. Each channel was low-pass filtered (<2 Hz) and a linear least-squared model fit the control signal (405 nm) to the dopamine-dependent signal (470 nm). Fluorescent signal (dF/F) was calculated as ([470 nm signal − fitted 405 nm signal]/[fitted 405 nm signal]). To measure slow fluctuations in fluorescent signal caused by injection of morphine or fentanyl, we first filtered the signal by calculating median fluorescence across a rolling 20 sec window, to remove transient events. This filtered signal was then averaged for the 30 minutes immediately before injection (baseline), and 30 mins immediately after morphine challenge or 5 mins immediately after fentanyl injection (i.e., area-under-the-curve averaged over time).

To measure spontaneous fluorescent transient events, we first corrected for photobleaching and other tonic signal fluctuations over time by subtracting the dF/F signal by the eighth percentile value over a rolling 20 sec window [4]. We identified spontaneous fluorescent transient events from this normalized signal as exceeding the median absolute mediation (MAD) by >2.91 deviations across a rolling 20 sec window [5]. For analysis of optogenetically-evoked signals, the onset and offset of the 595 nm LED were transmitted to the fiber photometry workstation as a TTL signal. The dLight signal on each stimulation trial was then normalized to a 2 sec baseline period immediately preceding stimulation, and the peak dLight signal during the 2 sec immediately following LED onset was averaged across the ten trials per frequency (0/5/10/20/40 Hz).

**Quantitative RT-PCR**

For quantitative RT-PCR, tissue was snap frozen on dry ice and stored at -80°C. RNA was isolated using the RNeasy Mini Kit (Qiagen) according to the manufacturer's instructions. All RNA samples had A260/A280 purity ratio ≥ 2. Reverse transcription was performed using Superscript III (Invitrogen). For each sample, duplicate cDNA preparations were set up. Mouse β-actin mRNA was used as the endogenous control. Quantitative RT-PCR using SYBR green (BioRad, Hercules, CA) was carried out with a Lightcycler 480 II (Roche) system with the following cycle parameters: 1 x (30 sec @ 95°C), 35 x (5 sec @ 95°C followed by 30 sec @ 60°C). Data were analyzed by comparing the C(t) values of the treatments tested using the ΔΔC(t) method. Expression values of target genes were first normalized to the expression value of β-actin. The median of each cDNA replicate reactions was used to quantify the relative target gene expression. Primers were designed in Primer3 and validated in BLAST and are listed in Table S1.

### RNA Sequencing

For RNA sequencing, tissue was stored in RNAlater (Qiagen) and total RNA was isolated using the RNeasy Mini Kit (Qiagen) as previously described [6]. Total eukaryotic RNA isolates were quantified using a fluorimetric RiboGreen assay. Total RNA integrity was assessed using capillary electrophoresis (Agilent BioAnalyzer 2100), generating an RNA Integrity Number (RIN) that was >8 for all samples. Total RNA samples were converted to Illumina sequencing libraries using the Truseq Stranded mRNA Sample Preparation Kit (Illumina, San Diego, CA, USA). Extracted mRNA was then oligo-dT purified using oligo-dT coated magnetic beads, fragmented and then reverse transcribed into cDNA. The cDNA was adenylated and then ligated to dual-indexed (barcoded) adapters. Truseq libraries were then sequenced 50-bp paired-end run on the Illumina HiSeq 2500, generating roughly 20 million paired-end reads per run.

Raw Illumina reads were cleaned of low quality bases, adapter contamination, and low complexity sequence with Trimmomatic [7]. Cleaned reads were aligned to the *Mus musculus* reference genome, version GRCm38 with HISAT2 [8], with a list of putative splice sites derived from the GTF annotation file. Resulting BAM files were sorted by read name, and counts were generated with HTSeq [9] with the unstranded option. A modification to the GTF file from Ensembl was performed to consider only protein coding genes for the counts. The expression counts matrix was filtered to remove genes with less than 150 total counts across all samples prior to differential expression testing. Nucleus accumbens and dorsal striatum samples were handled separately for all analyses.

### Normalization and Differential Expression Tests

For differential expression analysis, filtered expression counts were normalized and variance-stabilized with DESeq2 [10]. The variance-stabilized counts for all genes passing the expression filter were decomposed with principal components analysis (PCA) to identify potential broad patterns in gene expression differences among samples. Stabilized and filtered counts were also centered about 0 for each gene and used as input for hierarchical clustering. One sample from the nucleus accumbens was identified as an outlier based on both PCA and hierarchical clustering, and was removed from further analyses. Normalized and stabilized counts were exported for other analyses.

Differential expression testing was performed with normalized counts in DESeq2. No differences were observed between the saline-saline and saline-naloxone treatments (see Table S3), so these treatments were combined in a single "control" group for differential expression testing. Differentially expressed genes were identified among three comparisons: control vs. continuous, control vs. interrupted, and continuous vs. interrupted. A false discovery rate (q) of 0.05 was used for Benjamini-Hochberg multiple testing corrections [11], with a fold change threshold of 15%. For the purpose of comparison with prior literature [12], we also repeated this analysis using a less stringent cutoff of $p<0.05$ (see Figure S6).

**Pathway and Network Analyses**

Differentially expressed genes from DESeq were used to identify overrepresented pathways with Ingenuity Pathway Analysis (IPA) core expression analysis (QIAGEN Inc). Filtered, normalized, and stabilized counts from DESeq2 were scaled to mean 0 and unit variance was used as input for a weighted gene co-expression network analysis (WGCNA) [13,14]. Scale-free topology assumptions and soft power thresholds for network edges were checked with routines within the WGCNA package. Coexpression networks were built separately for each brain region, and then region-specific networks were merged to form a consensus network, merging modules with a Pearson correlation of at least 0.3 between their eigengenes. Eigengene values for each sample were exported. Tests for associations between module eigengene expression value and experimental treatment were performed with a one-way ANOVA. An eigengene-treatment association was considered as significant if its Bonferroni-adjusted P-value was less than 0.05, with the number of modules tested taken as the number of independent hypothesis tests.

**Statistical Analyses**

Similar numbers of male and female animals were used in all experiments, except for RNA sequencing, which was only conducted in males to minimize variability. Individual data points from males (filled circles) and females (open circles) are distinguished in figures. Sex was included as a variable in factorial ANOVA models analyzed using IBM SPSS Statistics v24, with repeated measures on within-subject factors, but sex effects were not significant unless noted otherwise. For main effects or interactions involving repeated measures, the Huynh-Feldt correction was applied to control for potential violations of the sphericity assumption. This correction reduces the degrees of freedom, resulting in non-integer values. Significant interactions were decomposed by analyzing simple effects (i.e., the effect of one variable at each level of the other variable). Significant main effects were analyzed using LSD post-hoc tests. Effect sizes are expressed as partial eta-squared ($\eta_p^2$) values. The Type I error rate was set to $\alpha=0.05$ (two-tailed) for all comparisons.

### SUPPLEMENTARY FIGURES & FIGURE LEGENDS

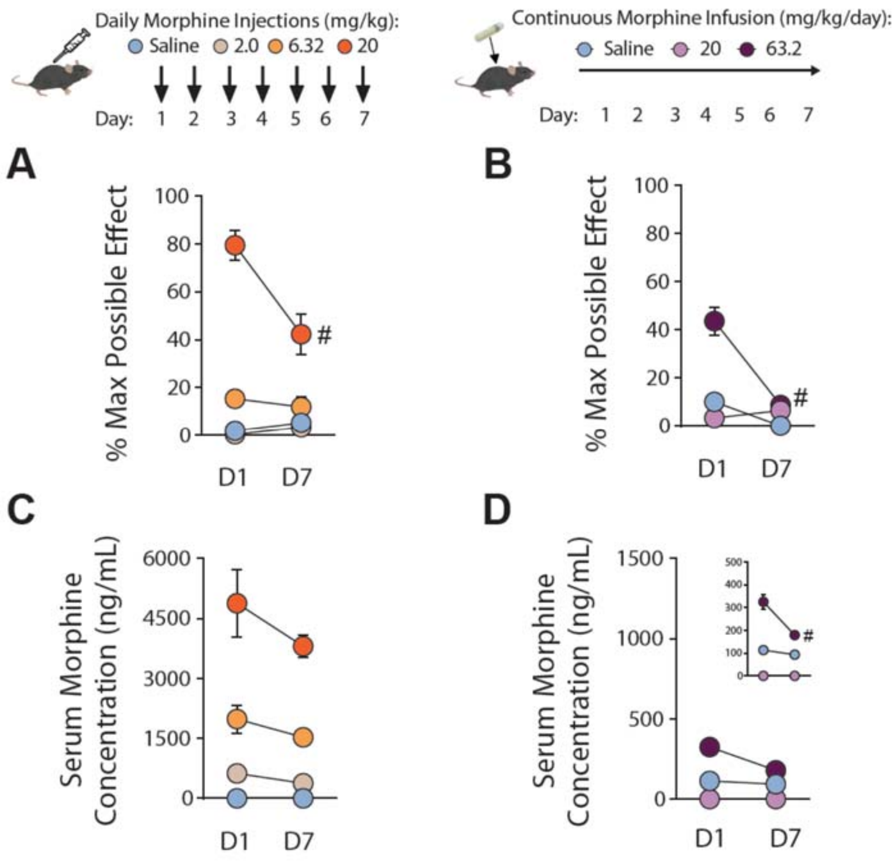


**Figure S1. Antinociception and pharmacokinetics after intermittent injection or continuous infusion of morphine. (A-B)** Thermal antinociception on the hot-plate increased in a dose-dependent fashion after morphine injections **(A)** (n=14/group) and infusion **(B)** (n=16/group). Tolerance was observed at the highest dose, for both injection and infusion. As noted in Supplemental Table S2, there was a significant Day x Dose x Sex interaction for morphine injections on the hot plate **(A)**. This interaction reflected a heightened antinociceptive response in female mice after 20mg/kg morphine injection (but not other doses), on Day 1 but not Day 7. **(C-D)** Serum morphine concentration increased in a dose-dependent fashion after injection **(C)** (n=5-6/group) or infusion **(D)** (n=8/group), with lower absolute levels after infusion (magnified in inset). All groups contained similar numbers of female and male mice; see Supplemental Table S2 for detailed statistical analyses. #p<0.05 for simple effect within group.

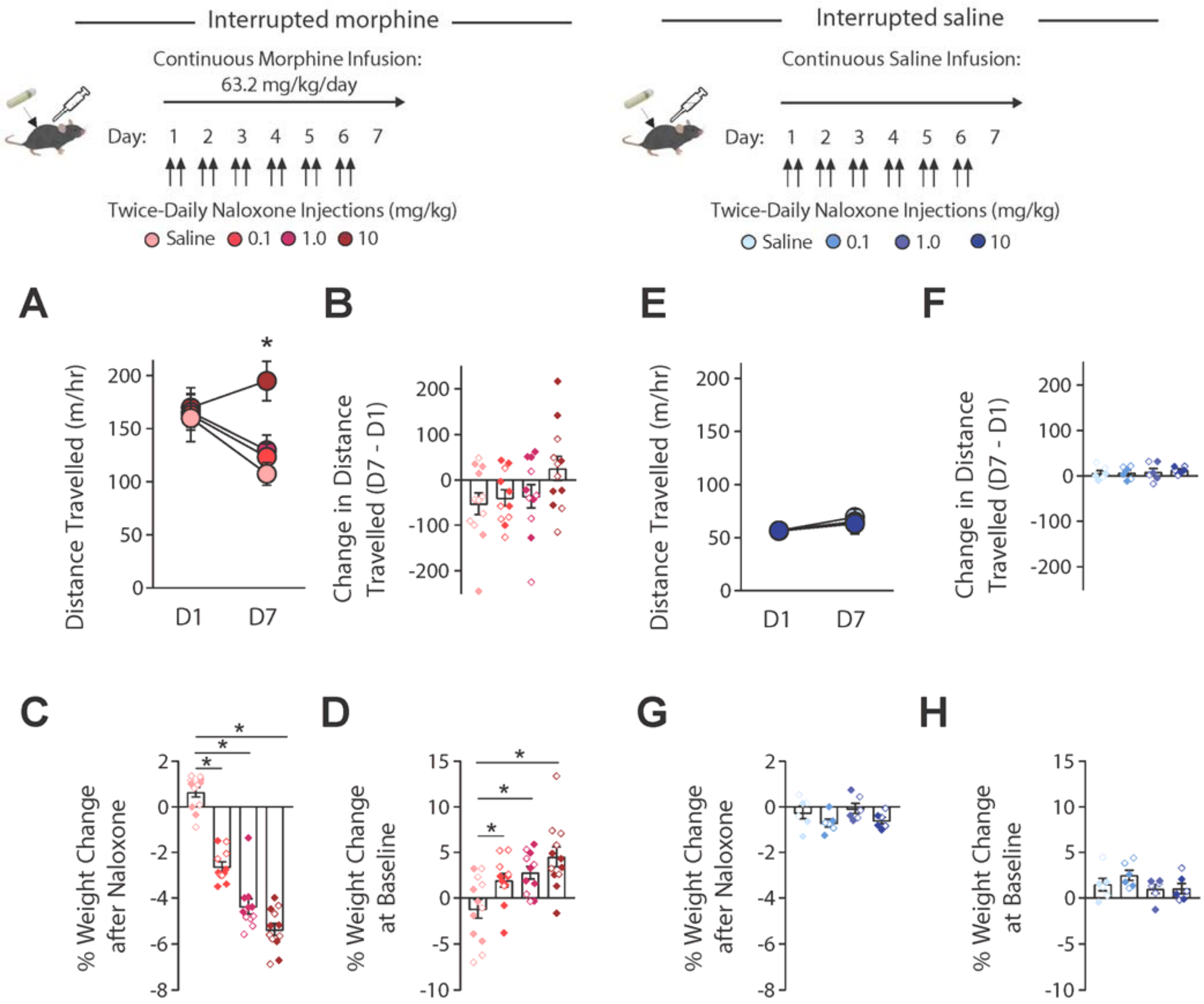


**Figure S2. Dose-response analysis for interruption of continuous morphine infusion with naloxone.** **(A)** Locomotor activity on the first (D1) and last (D7) day of exposure, showing gradual reversal of psychomotor tolerance with increasing doses of naloxone. *p<0.05, simple effect of Dose on D7. **(B)** Change in locomotor activity on D7 versus D1, depicted for individual mice at each dose. **(C)** Weight loss after the second injection of naloxone, compared to baseline weight before naloxone injection, and averaged across all six days of naloxone injections. *p<0.05 between groups, LSD post-hoc test. **(D)** Change in baseline weight on D7 versus D1, showing weight gain associated with naloxone treatment, due to an increase in food and water consumption (data not shown). *p<0.05 between groups, LSD post-hoc test. **(E-G)** No effects of naloxone in control groups implanted with saline pumps. All groups contained similar numbers of female mice (open symbols) and male mice (closed symbols); see Supplemental Table S2 for detailed statistical analyses.

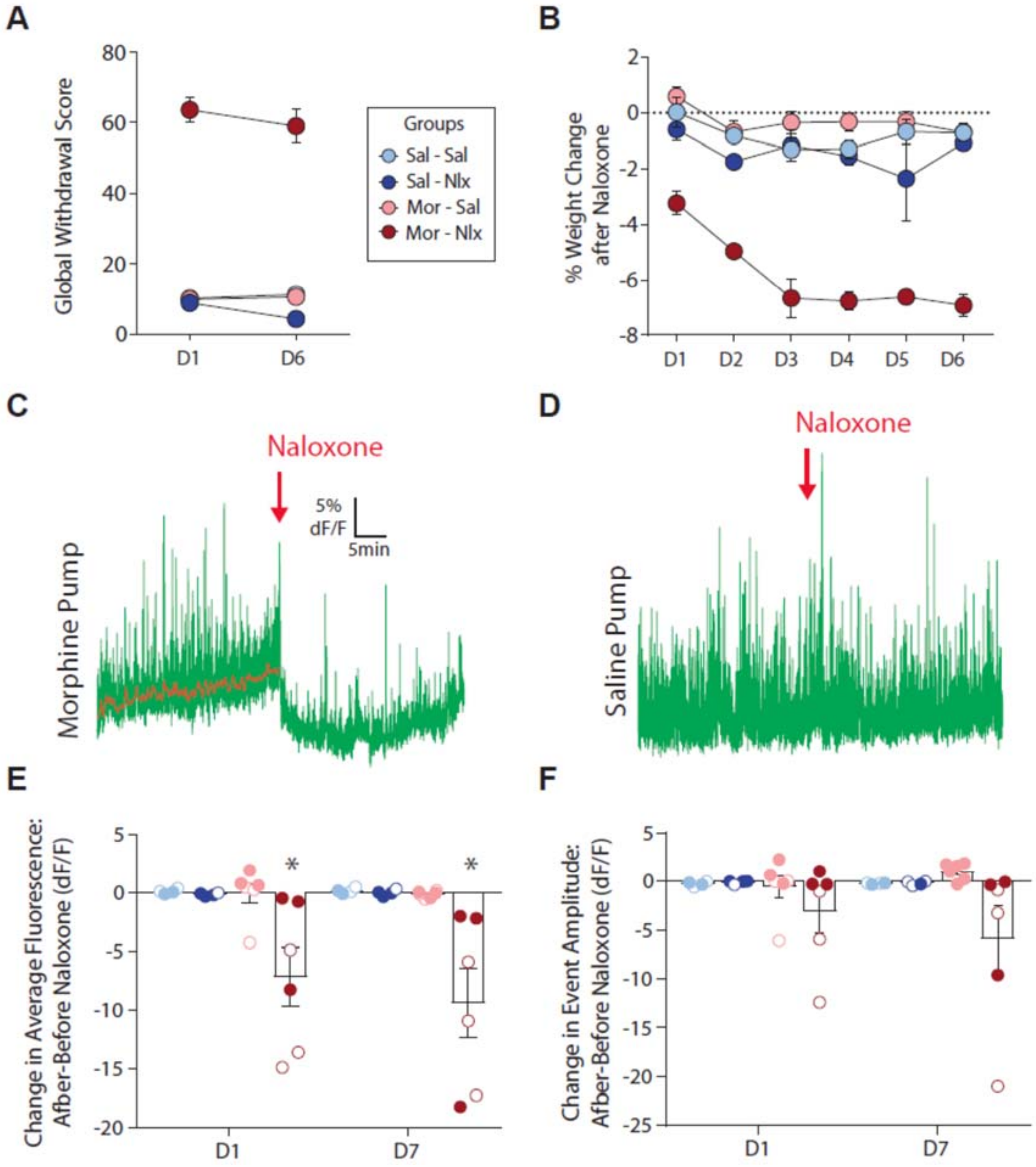


**Figure S3. Severity of naloxone-precipitated withdrawal across interrupted morphine treatment. (A)** Global withdrawal scores across days 1 and 6 of drug treatment, which were elevated in the morphine-naloxone group but did not change across days. **(B)** Weight loss following the second injection of naloxone on each day of treatment, compared with baseline weight before the first naloxone injection. **(C)** Example of decreasing dLight fluorescent signal following naloxone injection in a mouse implanted with a morphine pump. **(D)** Example of a stable dLight fluorescent signal following naloxone injection in a mouse implanted with a saline pump. **(E)** Change in dLight fluorescent signal following naloxone injection across days 1 and 7, depicted for individual mice. *p<0.05, decreased signal in morphine-naloxone group. **(F)** Change in spontaneous transient event amplitude across days 1 and 7, depicted for individual mice. All groups contained similar numbers of female mice (open symbols) and male mice (closed symbols); see Supplemental Table S2 for detailed statistical analyses.

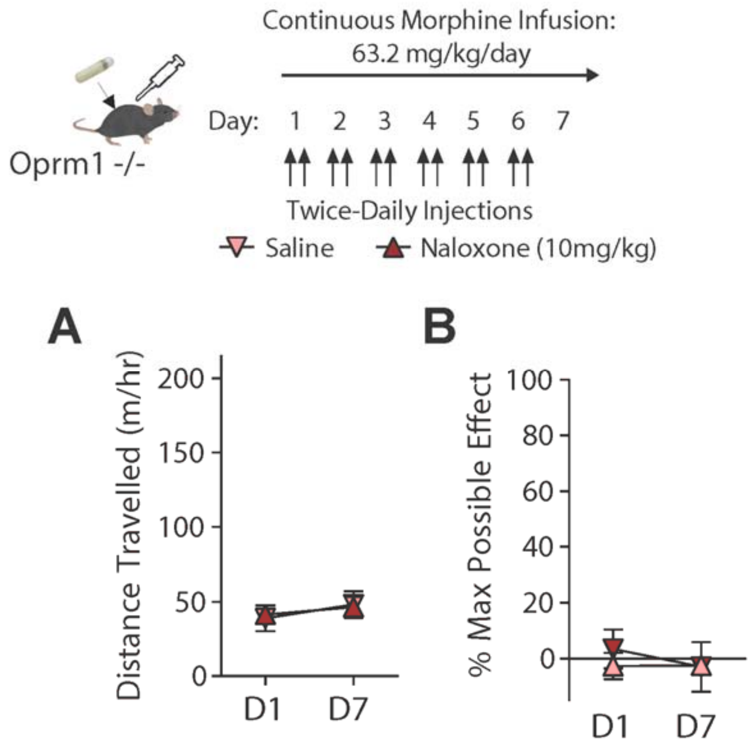


**Figure S4. Interrupted morphine treatment in mu opioid receptor knockout mice (Oprm1$^{-/-}$). (A)** Constitutive genetic deletion of the mu opioid receptor blocks acute psychomotor stimulation on the first day of morphine exposure (D1), as well as behavioral adaptations caused by chronic continuous or interrupted morphine exposure (D7). **(B)** Constitutive genetic deletion of the mu opioid receptor blocks thermal antinociception caused by morphine. Groups contained similar numbers of female and male mice; see Supplemental Table S2 for detailed statistical analyses.

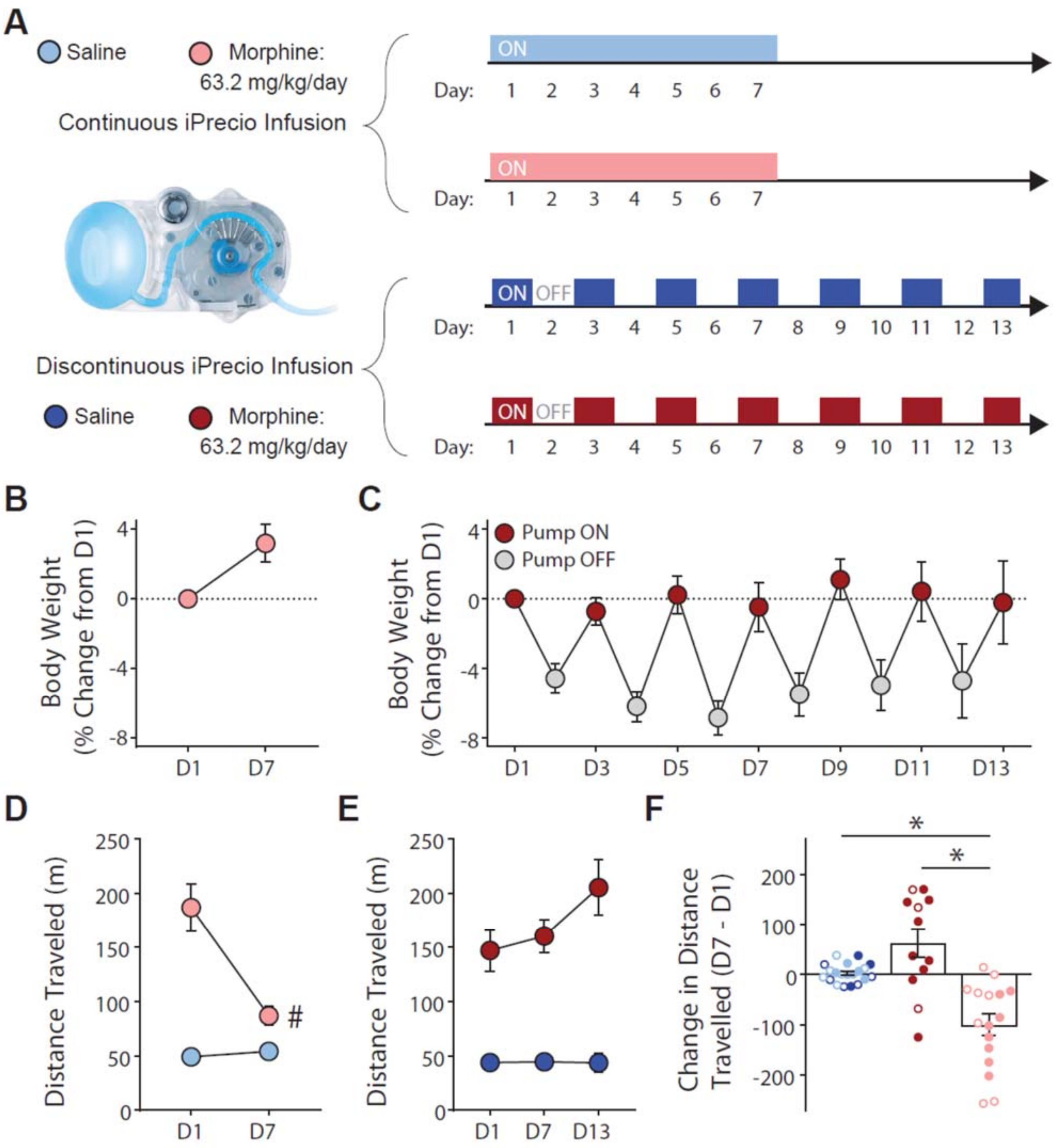


**Figure S5. Interruption of continuous morphine administration by spontaneous withdrawal. (A)** Image of miniaturized infusion pump (iPrecio SMP-300), and diagrams illustrating continuous administration over seven days (top), or discontinuous administration with alternating 24-hour periods of drug infusion and pump inactivity. Note that peak drug level (controlled by infusion rate) and total exposure (seven days at 63.2 mg/kg/day) are equivalent between continuous and discontinuous administration patterns. **(B-C)** Change in body weight during continuous morphine infusion **(B)** or discontinuous morphine infusion **(C)**, expressed as percent change from the first day of exposure (D1). Grey symbols in **(C)** indicate days when pump was inactive. **(D)** Locomotor activity after the first (D1) and last (D7) day of continuous exposure. **(E)** Locomotor activity after the first (D1), middle (D7), and last (D13) day of discontinuous exposure. **(F)** Change in locomotor activity on the last versus first day of treatment, with saline groups combined for clarity. All groups contained similar numbers of female mice (open symbols) and male mice (closed symbols); see Supplemental Table S2 for detailed statistical analyses. *$p<0.05$ between groups, LSD post-hoc test.

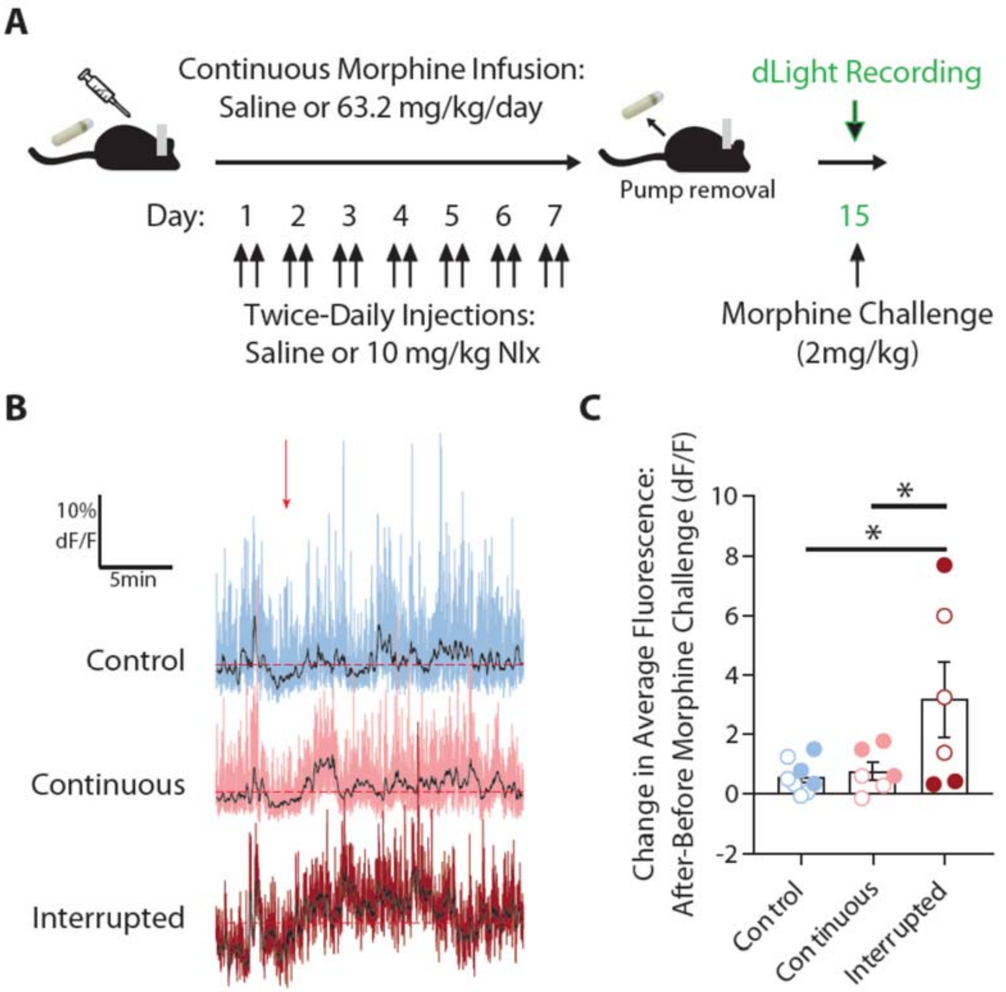


**Figure S6. Mesolimbic dopamine response to morphine challenge after interrupted morphine exposure. (A)** Experimental timeline for photometry recordings following morphine challenge. **(B)** Representative trace from each group in response to morphine challenge (2 mg/kg, red arrow). **(C)** Change in average fluorescent signal after challenge injection of morphine, compared to baseline signal before injection. All groups contained similar numbers of female mice (open symbols) and male mice (closed symbols); see Supplemental Table S2 for detailed statistical analyses. *p<0.05 between groups, LSD post-hoc test.

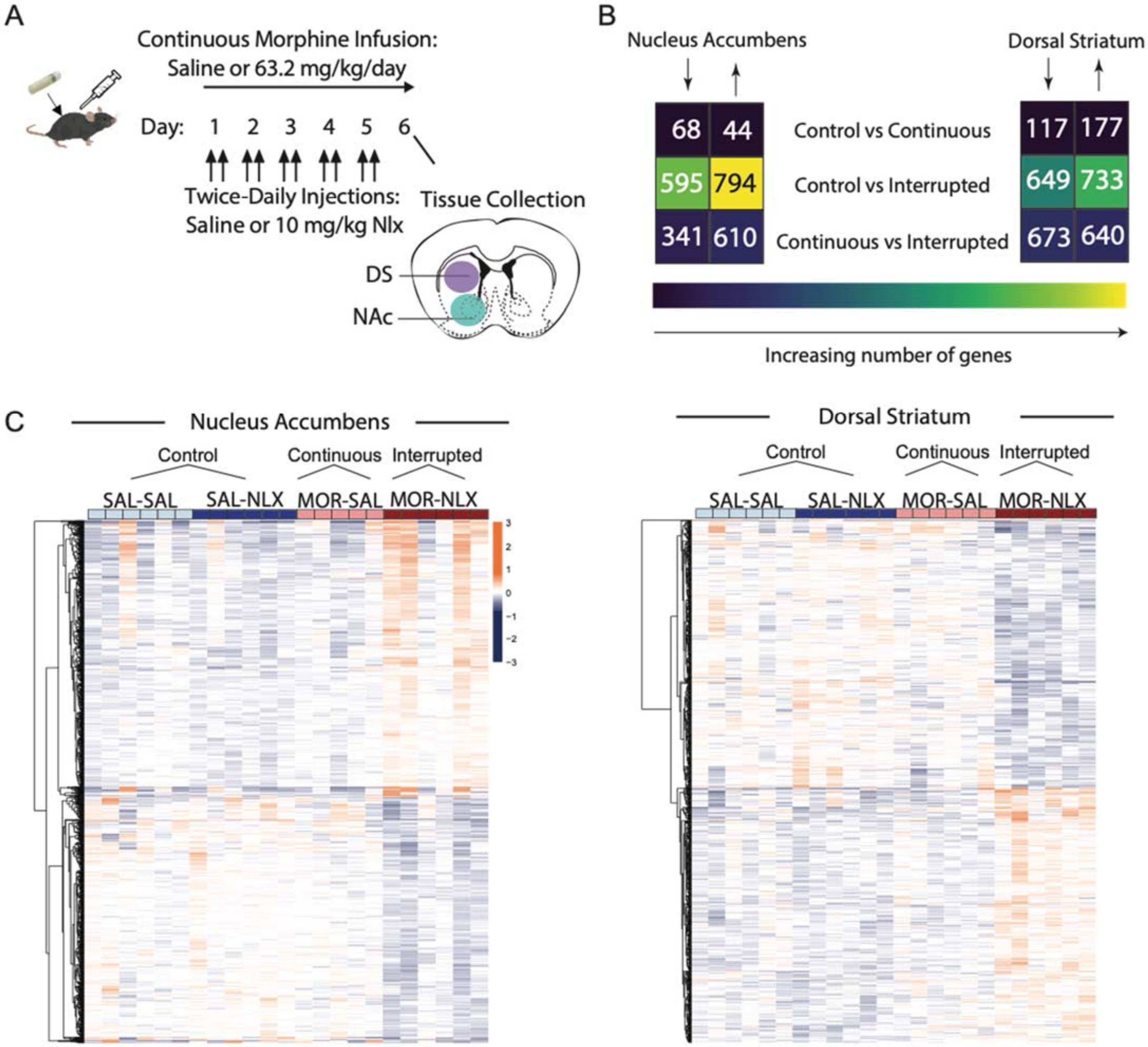


**Figure S7. Analysis of differential gene expression with unadjusted p-values. (A)** Microdissection of brain tissue for RNA sequencing from the nucleus accumbens (NAc) or dorsal striatum (DS), after six days of continuous or interrupted morphine exposure (n=5-6 male mice/group). **(B)** The number of differentially expressed genes (DEGs) that were significantly up- or down-regulated in each brain region; note that mice implanted with saline pumps have been combined to form a single control group. **(C)** Heat maps showing normalized level of DEGs for each individual sample.

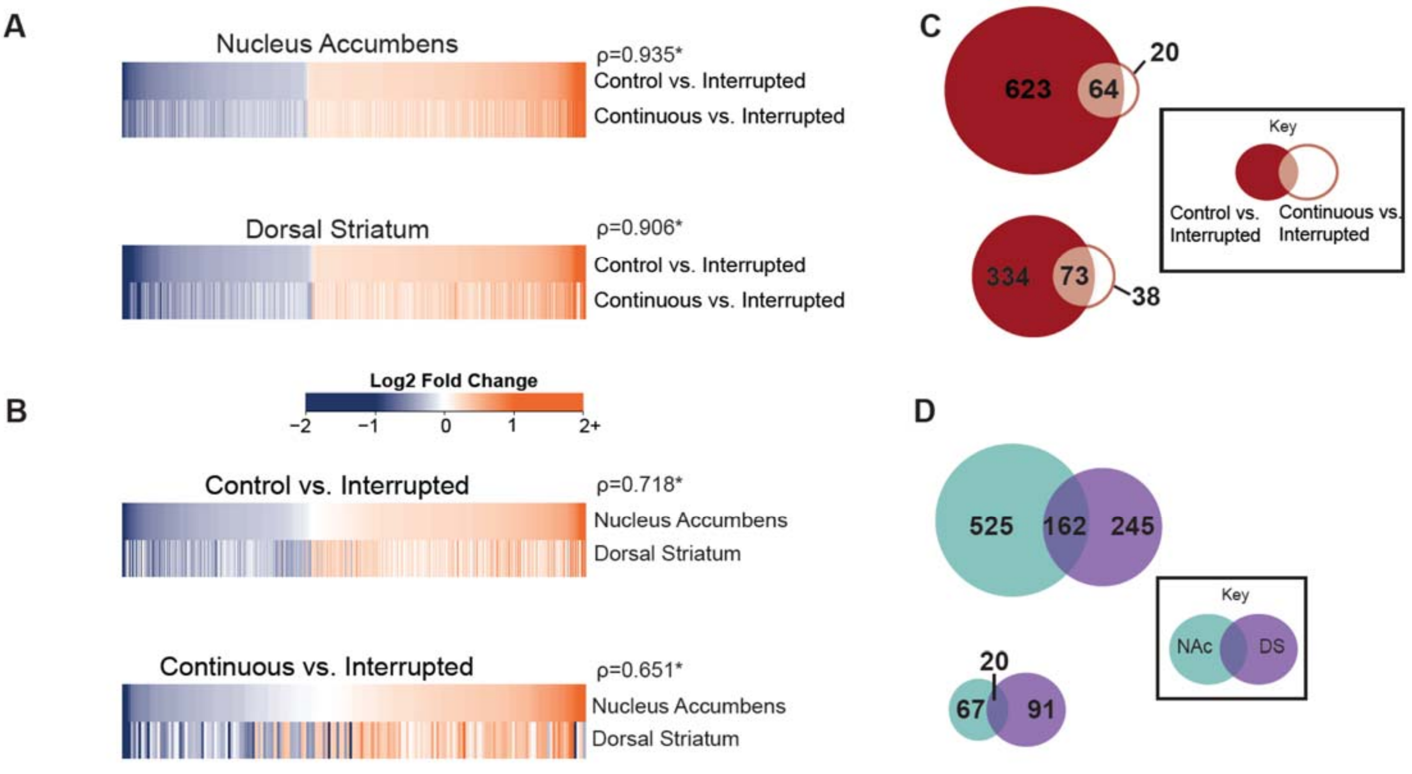


**Figure S8. Union of differential gene expression after continuous or interrupted morphine. (A-B)** Union heat maps showing aligned expression of genes in each brain region that were differentially expressed in comparisons between either Control vs. Interrupted or Continuous vs. Interrupted (**A**), or genes for each comparison that were differentially expressed in nucleus accumbens or dorsal striatum (**B**), aligned by fold change in the first comparison. **(C-D)** Venn diagrams showing the number of shared and unique DEGs for Control vs. Interrupted and Continuous vs. Interrupted comparisons in each brain region (**C**), or for nucleus accumbens and dorsal striatum in each comparison **(D)**.

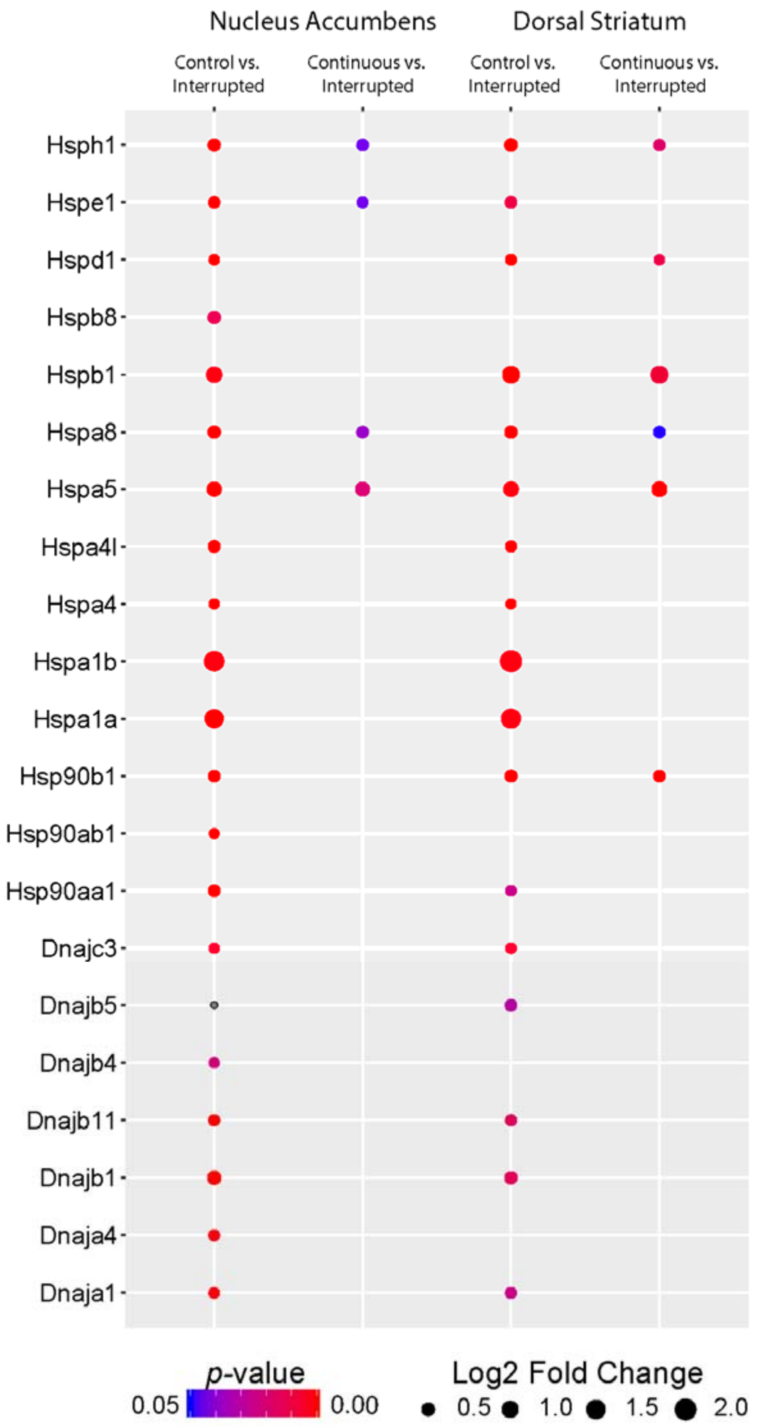


**Figure S9. Upregulated transcription of heat shock proteins after interrupted morphine exposure.** Columns represent individual comparisons between interrupted morphine and either control or continuous morphine for each brain region. Rows represent individual transcripts, with dot size representing fold change and dot color indicating statistical significance.

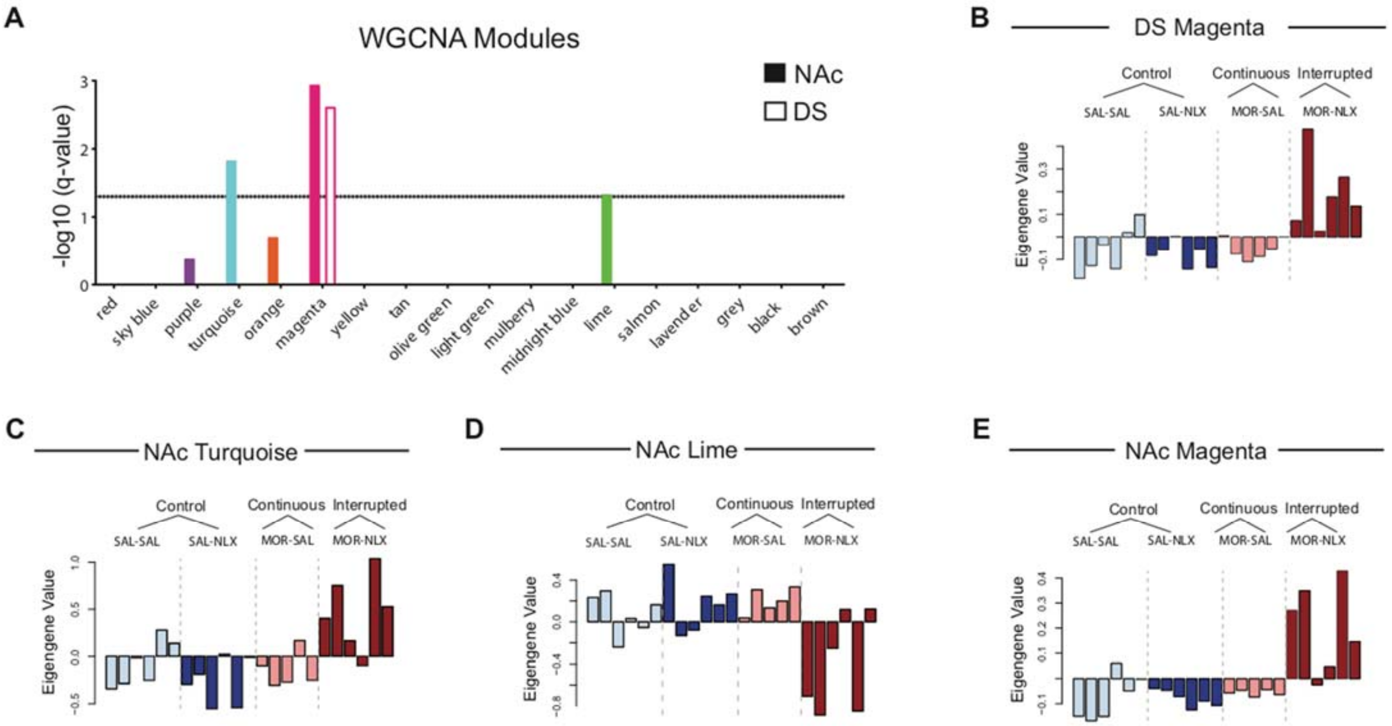


**Figure S10. Weighted gene co-expression network analysis. (A-E)** Consensus modules identified across the nucleus accumbens (NAc) and dorsal striatum (DS), with bars depicting Bonferroni-corrected p-value from a one-way ANOVA on eigengene values across treatment conditions **(A)**. Note that colors used to identify each module are arbitrary. The magnitude of gene regulation within each module can be summarized by an eigengene value calculated for each individual sample. The resulting eigengene values for individual samples from the dorsal striatum **(B)** and nucleus accumbens **(C-E)**, are shown for the magenta module **(B & E)**, turquoise module **(C)**, and lime module **(D)**. In all three modules, large eigengene values were only observed after interrupted morphine, while eigengene values after continuous morphine group were similar to control values.

**Table S1:** List of primers used for quantitative RT-PCR.

| Gene Name | Symbol | Forward oligonucleotide | Reverse oligonucleotide |
|---|---|---|---|
| beta-actin | β-actin | GACGGCCAGGTCATCACTAT | CCACCGATCCACACAGAGTA |
| RNA binding motif protein 3 | Rbm3 | CCTTCACAAACCCAGAGCAT | TAGACCGCCCATACCCATA |
| Cold-inducible RNA-binding protein | Cirbp | CTTAGGAAGCTTGGGTGTGT | CGTCCTTAGCGTCATCGATATT |
| Heat shock protein family B (Small) member 1 | Hspb1 | ACCCTAGTGTCCTCTTCCCT | GCGCACAGATTGACAGAGAG |
| Heat shock protein family A (Hsp70) member 5 | Hspa5 | CCCCAACTGGTGAAGAGGAT | CCCCAAGACATGTGAGCAAC |

**Table S2**. Comprehensive reporting of all main effects and interactions from ANOVA models, including those involving sex. Effect sizes are expressed as partial eta-squared ($\eta_p^2$) values.

| Figure | Factor | F-value | p-value | $\eta_p^2$ |
|---|---|---|---|---|
| Figure 1B: Distance travelled during intermittent exposure | Dose (between) | $F_{4,62} = 94.91$ | <0.001* | 0.860 |
| | Sex (between) | $F_{1,62} < 1$ | | |
| | Dose x Sex | $F_{4,62} < 1$ | | |
| | Day (within) | $F_{1,62} = 188.13$ | <0.001* | 0.752 |
| | Day x Dose | $F_{4,62} = 46.84$ | <0.001* | 0.751 |
| | Day x Sex | $F_{1,62} < 1$ | | |
| | Day x Dose x Sex | $F_{4,62} < 1$ | | |
| Figure 1D: Distance travelled after challenge injections following intermittent pretreatment | Pretreatment Dose (between) | $F_{4,62} = 14.32$ | <0.001* | 0.480 |
| | Sex (between) | $F_{1,62} < 1$ | | |
| | Pretreatment Dose x Sex | $F_{4,62} < 1$ | | |
| | Challenge Dose (within) | $F_{2.48,153.75} = 544.67$ | <0.001* | 0.898 |
| | Challenge Dose x Pretreatment Dose | $F_{9.92, 153.75} = 8.55$ | <0.001* | 0.356 |
| | Challenge Dose x Sex | $F_{2.48, 153.75} < 1$ | | |
| | Challenge Dose x Pretreatment Dose x Sex | $F_{9.92, 153.75} < 1$ | | |
| Figure 1F: Distance travelled during continuous exposure | Dose (between) | $F_{2,42} = 40.83$ | <0.001* | 0.660 |
| | Sex (between) | $F_{2,42} = 1.06$ | 0.309 | 0.025 |
| | Dose x Sex | $F_{2,42} = 1.12$ | 0.334 | 0.051 |
| | Day (within) | $F_{1,42} = 7.71$ | 0.008* | 0.155 |
| | Day x Dose | $F_{2,42} = 8.94$ | 0.001* | 0.299 |
| | Day x Sex | $F_{1,42} < 1$ | | |
| | Day x Dose x Sex | $F_{2,42} < 1$ | | |
| Figure 1H: Distance travelled after challenge injections following continuous pretreatment | Pretreatment Dose (between) | $F_{2,18} = 1.48$ | 0.253 | 0.141 |
| | Sex (between) | $F_{1,18} < 1$ | | |
| | Pretreatment Dose x Sex | $F_{2,18} < 1$ | | |
| | Challenge Dose (within) | $F_{2.87,51.73} = 217.32$ | <0.001* | 0.924 |
| | Challenge Dose x Pretreatment Dose | $F_{5.75,51.73} = 2.00$ | 0.085 | 0.182 |
| | Challenge Dose x Sex | $F_{2.87,51.73} < 1$ | | |
| | Challenge Dose x Pretreatment Dose x Sex | $F_{5.75,51.73} < 1$ | | |

| Figure | Factor | F-value | p-value | $\eta_p^2$ |
|---|---|---|---|---|
| Figure 2B: Distance travelled during interrupted exposure | Morphine (between) | $F_{1,86}$ = 171.26 | <0.001* | 0.904 |
| | Naloxone (between) | $F_{1,86}$ = 9.03 | 0.003 | 0.095 |
| | Morphine x Naloxone | $F_{1,86}$ = 9.02 | 0.004 | 0.095 |
| | Sex (between) | $F_{1,86}$ < 1 | | |
| | Sex x Morphine | $F_{1,86}$ < 1 | | |
| | Sex x Naloxone | $F_{1,86}$ < 1 | | |
| | Sex x Morphine x Naloxone | $F_{1,86}$ < 1 | | |
| | Day (Within) | $F_{1,86}$ < 1 | | |
| | Day x Morphine | $F_{1,86}$ < 1 | | |
| | Day x Naloxone | $F_{1,86}$ = 10.14 | 0.002* | 0.105 |
| | Day x Sex | $F_{1,86}$ = 1.18 | 0.281 | 0.014 |
| | Day x Morphine x Naloxone | $F_{1,86}$ = 10.32 | 0.002* | 0.107 |
| | Day x Morphine x Sex | $F_{1,86}$ = 1.30 | 0.257 | 0.015 |
| | Day x Naloxone x Sex | $F_{1,86}$ < 1 | | |
| | Day x Morphine x Naloxone x Sex | $F_{1,86}$ < 1 | | |
| Figure 2D: Serum morphine levels during interrupted exposure | Naloxone (between) | $F_{1,6}$ < 1 | | |
| | Sex (between) | $F_{1,6}$ < 1 | | |
| | Naloxone x Sex | $F_{1,6}$ < 1 | | |
| | Day (within) | $F_{1,6}$ = 27.90 | 0.002* | 0.823 |
| | Day x Naloxone | $F_{1,6}$ = 3.78 | 0.100 | 0.386 |
| | Day x Sex | $F_{1,6}$ = 2.61 | 0.157 | 0.303 |
| | Day x Sex x Naloxone | $F_{1,6}$ = 2.12 | 0.196 | 0.261 |
| Figure 2E: Thermal antinociception during interrupted exposure | Morphine (between) | $F_{1,38}$ = 21.17 | <0.001* | 0.358 |
| | Naloxone (between) | $F_{1,38}$ < 1 | | |
| | Morphine x Naloxone | $F_{1,38}$ < 1 | | |
| | Sex (between) | $F_{1,38}$ < 1 | | |
| | Sex x Morphine | $F_{1,38}$ < 1 | | |
| | Sex x Naloxone | $F_{1,38}$ < 1 | | |
| | Sex x Morphine x Naloxone | $F_{1,38}$ < 1 | | |
| | Day (within) | $F_{1,38}$ = 10.25 | 0.003* | 0.212 |
| | Day x Morphine | $F_{1,38}$ = 11.83 | 0.001* | 0.237 |
| | Day x Naloxone | $F_{1,38}$ = 1.64 | 0.208 | 0.041 |
| | Day x Sex | $F_{1,38}$ = 2.10 | 0.156 | 0.052 |
| | Day x Morphine x Sex | $F_{1,38}$ < 1 | | |
| | Day x Naloxone x Sex | $F_{1,38}$ < 1 | | |
| | Day x Morphine x Naloxone | $F_{1,38}$ < 1 | | |
| | Day x Morphine x Naloxone x Sex | $F_{1,38}$ < 1 | | |

| Figure | Factor | F-value | p-value | $\eta_p^2$ |
|---|---|---|---|---|
| Figure 2F: Distance travelled after saline challenge at 24 hours of withdrawal | Morphine (between) | $F_{1,64} = 6.90$ | 0.011* | 0.097 |
| | Naloxone (between) | $F_{1,64} < 1$ | | |
| | Morphine x Naloxone | $F_{1,64} = 1.36$ | 0.247 | 0.021 |
| | Sex (between) | $F_{1,64} = 1.84$ | 0.180 | 0.028 |
| | Sex x Morphine | $F_{1,64} = 2.08$ | 0.154 | 0.032 |
| | Sex x Naloxone | $F_{1,64} < 1$ | | |
| | Sex x Morphine x Naloxone | $F_{1,64} = 1.12$ | 0.293 | 0.017 |
| Figure 2G: Distance travelled after morphine challenge at 48 hours of withdrawal | Morphine (between) | $F_{1,64} < 1$ | | |
| | Naloxone (between) | $F_{1,64} = 15.34$ | <0.001* | 0.193 |
| | Morphine x Naloxone | $F_{1,64} = 13.37$ | 0.001* | 0.173 |
| | Sex (between) | $F_{1,64} = 1.74$ | 0.192 | 0.026 |
| | Sex x Morphine | $F_{1,64} < 1$ | | |
| | Sex x Naloxone | $F_{1,64} < 1$ | | |
| | Sex x Morphine x Naloxone | $F_{1,64} = 2.28$ | 0.136 | 0.034 |
| Figure 2H: Distance travelled after morphine challenges during extended withdrawal (naloxone groups) | Morphine (between) | $F_{1,17} = 13.87$ | 0.002* | 0.449 |
| | Sex (between) | $F_{1,17} < 1$ | | |
| | Morphine x Sex | $F_{1,17} < 1$ | | |
| | Day (within) | $F_{10.23,174.00} = 5.13$ | <0.001* | 0.232 |
| | Day x Morphine | $F_{10.23,174.00} = 1.81$ | 0.060 | 0.096 |
| | Day x Sex | $F_{10.23,174.00} < 1$ | | |
| | Day x Morphine x Sex | $F_{10.23,174.00} < 1$ | | |
| Figure 2I: Distance travelled after morphine challenges during extended withdrawal (saline groups) | Morphine (between) | $F_{1,18} = 1.55$ | 0.229 | 0.079 |
| | Sex (between) | $F_{1,18} = 2.32$ | 0.145 | 0.114 |
| | Morphine x Sex | $F_{1,18} < 1$ | | |
| | Day (within) | $F_{8.33,149.96} = 5.96$ | <0.001* | 0.237 |
| | Day x Morphine | $F_{8.33,149.96} = 1.40$ | 0.198 | 0.072 |
| | Day x Sex | $F_{8.33,149.96} < 1$ | | |
| | Day x Morphine x Sex | $F_{8.33,149.96} < 1$ | | |
| Figure 3H: Event Amplitude on Days 0, 1 and 7 | Group (between) | $F_{2,14} = 2.09$ | 0.160 | 0.231 |
| | Sex (between) | $F_{1,14} = 2.09$ | 0.170 | 0.130 |
| | Group x Sex | $F_{2,14} < 1$ | | |
| | Day (within) | $F_{2,28} = 6.00$ | 0.007* | 0.300 |
| | Day x Group | $F_{4,28} = 5.20$ | 0.006* | 0.397 |
| | Day x Sex | $F_{2,28} = 1.30$ | 0.288 | 0.085 |
| | Day x Group x Sex | $F_{4,28} = 1.31$ | 0.290 | 0.158 |

| Figure | Factor | F-value | p-value | $\eta_p^2$ |
|---|---|---|---|---|
| Figure 3I: Change in Event Amplitude: Day 7 - Day 1 | Group (between) | $F_{2,14} = 2.71$ | 0.101 | 0.279 |
| | Sex (between) | $F_{2,14} < 1$ | | |
| | Group x Sex | $F_{2,14} = 1.70$ | 0.218 | 0.196 |
| Figure 3M: Change in Average Fluorescence: Challenge – Acute Fentanyl | Group (between) | $F_{2,8} = 6.23$ | 0.023* | 0.609 |
| | Sex (between) | $F_{1,8} < 1$ | | |
| | Group x Sex | $F_{2,8} = 1.65$ | 0.252 | 0.292 |
| Figure 4I & K: Rbm3 mRNA expression (Note: "Subregion" refers to nucleus accumbens or dorsal striatum) | Group (between) | $F_{2,35} = 16.84$ | <0.001* | .490 |
| | Sex (between) | $F_{1,35} =1.44$ | 0.238 | 0.040 |
| | Group x Sex | $F_{2,35} < 1$ | | |
| | Subregion (within) | $F_{1,35} < 1$ | | |
| | Subregion x Group | $F_{2,35} < 1$ | | |
| | Subregion x Sex | $F_{1,35} < 1$ | | |
| | Subregion x Group x Sex | $F_{2,35} < 1$ | | |
| Figure 4J & L: Cirbp mRNA expression (Note: "Subregion" refers to nucleus accumbens or dorsal striatum) | Group (between) | $F_{2,35} = 12.29$ | <0.001* | 0.964 |
| | Sex (between) | $F_{1,35} < 1$ | 0.238 | 0.040 |
| | Group x Sex | $F_{2,35} = 1.60$ | 0.216 | 0.084 |
| | Subregion (within) | $F_{1,35} = 5.98$ | 0.020* | 0.146 |
| | Subregion x Group | $F_{2,35} = 2.06$ | 0.143 | 0.105 |
| | Subregion x Sex | $F_{1,35} < 1$ | | |
| | Subregion x Group x Sex | $F_{2,35} < 1$ | | |
| Figure 4M & O: Hspb1 mRNA expression (Note: "Subregion" refers to nucleus accumbens or dorsal striatum) | Group (between) | $F_{2,32} = 3.32$ | 0.049* | 0.172 |
| | Sex (between) | $F_{1,32} < 1$ | | |
| | Group x Sex | $F_{2,32} = 2.58$ | 0.091 | 0.139 |
| | Subregion (within) | $F_{1,32} < 1$ | | |
| | Subregion x Group | $F_{1,32} < 1$ | | |
| | Subregion x Sex | $F_{1,32} < 1$ | | |
| | Subregion x Group x Sex | $F_{1,32} < 1$ | | |
| Figure 4N & P: Hspb1 mRNA expression (Note: "Subregion" refers to nucleus accumbens or dorsal striatum) | Group (between) | $F_{2,33} = 3.65$ | 0.037* | 0.181 |
| | Sex (between) | $F_{1,33} < 1$ | | |
| | Group x Sex | $F_{2,33} = 2.89$ | 0.070 | 0.149 |
| | Subregion (within) | $F_{2,33} = 3.581$ | 0.067 | 0.098 |
| | Subregion x Group | $F_{2,33} = 2.041$ | 0.146 | 0.110 |
| | Subregion x Sex | $F_{1,33} < 1$ | | |
| | Subregion x Group x Sex | $F_{1,33} < 1$ | | |

| Figure | Factor | F-value | p-value | $\eta_p^2$ |
|---|---|---|---|---|
| Figure S1A: Serum morphine level during intermittent exposure | Dose (between) | $F_{2,10} = 29.11$ | <0.001* | 0.866 |
| | Sex (between) | $F_{1,10} = 1.22$ | 0.298 | 0.119 |
| | Dose x Sex | $F_{2,10} < 1$ | | |
| | Day (within) | $F_{1,9} = 5.28$ | 0.047* | 0.370 |
| | Day x Dose | $F_{2,9} < 1$ | | |
| | Day x Sex | $F_{1,9} < 1$ | | |
| | Day x Dose x Sex | $F_{2,9} < 1$ | | |
| Figure S1A: Serum morphine level during intermittent exposure | Dose (between) | $F_{2,10} = 29.11$ | <0.001* | 0.866 |
| | Sex (between) | $F_{1,10} = 1.22$ | 0.298 | 0.119 |
| | Dose x Sex | $F_{2,10} < 1$ | | |
| | Day (within) | $F_{1,9} = 5.28$ | 0.047* | 0.370 |
| | Day x Dose | $F_{2,9} < 1$ | | |
| | Day x Sex | $F_{1,9} < 1$ | | |
| | Day x Dose x Sex | $F_{2,9} < 1$ | | |
| Figure S1B: Serum morphine level during continuous exposure | Dose (between) | $F_{1,10} = 81.17$ | <0.001* | 0.890 |
| | Sex (between) | $F_{1,10} = 1.03$ | 0.334 | 0.093 |
| | Dose x Sex | $F_{1,10} < 1$ | | |
| | Day (within) | $F_{1,10} = 9.18$ | 0.013* | 0.478 |
| | Day x Dose | $F_{1,10} = 7.04$ | 0.024* | 0.413 |
| | Day x Sex | $F_{1,10} < 1$ | | |
| | Day x Dose x Sex | $F_{1,10} < 1$ | | |
| Figure S1C: Thermal antinociception during intermittent exposure | Dose (between) | $F_{3,40} = 59.89$ | <0.001* | 0.720 |
| | Sex (between) | $F_{1,40} = 3.19$ | 0.082 | 0.074 |
| | Dose x Sex | $F_{3,40} = 2.25$ | 0.097 | 0.145 |
| | Day (within) | $F_{1,40} = 9.25$ | 0.004* | 0.188 |
| | Day x Dose | $F_{3,40} = 14.33$ | <0.001* | 0.518 |
| | Day x Sex | $F_{1,40} < 1$ | | |
| | Day x Dose x Sex | $F_{3,40} = 3.74$ | 0.018* | 0.219 |
| Figure S1D: Thermal antinociception during continuous exposure | Dose (between) | $F_{2,42} = 30.16$ | <0.001* | 0.589 |
| | Sex (between) | $F_{1,42} = 2.88$ | 0.097 | 0.064 |
| | Dose x Sex | $F_{2,42} < 1$ | | |
| | Day (within) | $F_{1,42} = 30.25$ | <0.001* | 0.419 |
| | Day x Dose | $F_{2,42} = 19.50$ | <0.001* | 0.482 |
| | Day x Sex | $F_{1,42} < 1$ | | |
| | Day x Dose x Sex | $F_{2,42} < 1$ | | |

| Figure | Factor | F-value | p-value | $\eta_p^2$ |
|---|---|---|---|---|
| Figure S2A: Distance travelled during interrupted morphine exposure | Dose (between) | $F_{3,38} = 3.39$ | 0.028* | 0.211 |
| | Sex (between) | $F_{1,38} < 1$ | | |
| | Dose x Sex | $F_{1,38} < 1$ | | |
| | Day (within) | $F_{1,38} = 5.05$ | 0.031* | 0.117 |
| | Day x Dose | $F_{3,38} = 2.23$ | 0.100 | 0.150 |
| | Day x Sex | $F_{1,38} = 2.14$ | 0.152 | 0.053 |
| | Day x Dose x Sex | $F_{3,38} < 1$ | | |
| Figure S2C: Weight loss after naloxone injections | Dose (between) | $F_{3,38} = 118.32$ | <0.001* | 0.903 |
| | Sex (between) | $F_{1,38} = 1.33$ | 0.256 | 0.034 |
| | Dose x Sex | $F_{3,38} = 2.02$ | 0.127 | 0.138 |
| Figure S2D: Weight gain before naloxone injections | Dose (between) | $F_{3,38} = 7.61$ | <0.001* | 0.375 |
| | Sex (between) | $F_{1,38} = 3.75$ | 0.060 | 0.090 |
| | Dose x Sex | $F_{3,38} = 1.56$ | 0.215 | 0.110 |
| Figure S2E: Distance travelled during interrupted saline exposure | Dose (between) | $F_{3,16} < 1$ | | |
| | Sex (between) | $F_{1,16} < 1$ | | |
| | Dose x Sex | $F_{3,16} < 1$ | | |
| | Day (Within) | $F_{1,16} = 7.90$ | 0.013* | 0.331 |
| | Day x Dose | $F_{3,16} < 1$ | | |
| | Day x Sex | $F_{1,16} < 1$ | | |
| | Day x Dose x Sex | $F_{3,16} < 1$ | | |
| Figure S2G: Weight loss after naloxone injections | Dose (between) | $F_{3,16} = 1.93$ | 0.165 | 0.266 |
| | Sex (between) | $F_{1,16} < 1$ | | |
| | Sex x Dose | $F_{3,16} < 1$ | | |
| Figure S2H: Weight gain before naloxone injections | Dose (Between) | $F_{3,16} = 1.32$ | 0.304 | 0.198 |
| | Sex (Between) | $F_{1,16} = 1.51$ | 0.236 | 0.086 |
| | Sex x Dose | $F_{3,16} < 1$ | | |

| Figure | Factor | F-value | p-value | $\eta_p^2$ |
|---|---|---|---|---|
| Figure S3A: Global withdrawal score across days | Morphine (between) | $F_{1,40} = 148.9$ | <0.001* | 0.909 |
| | Naloxone (between) | $F_{1,40} = 153.1$ | <0.001* | 0.788 |
| | Morphine x Naloxone | $F_{1,40} = 110.2$ | <0.001* | 0.734 |
| | Sex (between) | $F_{1,40} < 1$ | | |
| | Sex x Morphine | $F_{1,40} < 1$ | | |
| | Sex x Naloxone | $F_{1,40} < 1$ | | |
| | Sex x Morphine x Naloxone | $F_{1,40} < 1$ | | |
| | Day (within) | $F_{1,40} = 1.56$ | 0.219 | 0.038 |
| | Day x Morphine | $F_{1,40} < 1$ | | |
| | Day x Naloxone | $F_{1,40} < 1$ | | |
| | Day x Sex | $F_{1,40} = 1.63$ | 0.209 | 0.039 |
| | Day x Morphine x Sex | $F_{1,40} < 1$ | | |
| | Day x Naloxone x Sex | $F_{1,40} < 1$ | | |
| | Day x Morphine x Naloxone | $F_{1,40} = 3.62$ | 0.064 | 0.083 |
| | Day x Morphine x Naloxone x Sex | $F_{1,40} < 1$ | | |
| Figure S3B: Weight change after naloxone across days | Morphine (between) | $F_{1,40} = 85.35$ | <0.001* | 0.681 |
| | Naloxone (between) | $F_{1,40} = 210.25$ | <0.001* | 0.840 |
| | Morphine x Naloxone | $F_{1,40} = 134.51$ | <0.001* | 0.771 |
| | Sex (between) | $F_{1,40} < 1$ | | |
| | Sex x Morphine | $F_{1,40} < 1$ | | |
| | Sex x Naloxone | $F_{1,40} = 5.10$ | 0.029* | 0.113 |
| | Sex x Morphine x Naloxone | $F_{1,40} < 1$ | | |
| | Day (within) | $F_{3.47,138.93} = 7.71$ | <0.001* | 0.162 |
| | Day x Morphine | $F_{3.47,138.93} = 1.87$ | 0.128 | 0.045 |
| | Day x Naloxone | $F_{3.47,138.93} = 1.75$ | 0.152 | 0.042 |
| | Day x Sex | $F_{3.47,138.93} = 1.54$ | 0.201 | 0.037 |
| | Day x Morphine x Sex | $F_{3.47,138.93} = 1.99$ | 0.108 | 0.047 |
| | Day x Naloxone x Sex | $F_{3.47,138.93} = 1.50$ | 0.212 | 0.036 |
| | Day x Morphine x Naloxone | $F_{3.47,138.93} = 2.37$ | 0.065 | 0.056 |
| | Day x Morphine x Naloxone x Sex | $F_{3.47,138.93} = 1.19$ | 0.319 | 0.029 |

| Figure | Factor | F-value | p-value | $\eta_p^2$ |
|---|---|---|---|---|
| Figure S3E: Change in average fluorescence: after-before naloxone injection | Morphine (between) | $F_{1,12} = 5.92$ | 0.032* | 0.331 |
| | Naloxone (between) | $F_{1,12} = 5.83$ | 0.033* | 0.327 |
| | Morphine x Naloxone | $F_{1,12} = 5.30$ | 0.040* | 0.306 |
| | Sex (between) | $F_{1,12} < 1$ | | |
| | Sex x Morphine | $F_{1,12} = 1.10$ | 0.316 | 0.084 |
| | Sex x Naloxone | $F_{1,12} < 1$ | | |
| | Sex x Morphine x Naloxone | $F_{1,12} < 1$ | | |
| | Day (within) | $F_{1,12} < 1$ | | |
| | Day x Morphine | $F_{1,12} = 1.19$ | 0.297 | 0.090 |
| | Day x Naloxone | $F_{1,12} < 1$ | | |
| | Day x Sex | $F_{1,12} = 1.80$ | 0.204 | 0.131 |
| | Day x Morphine x Sex | $F_{1,12} = 1.80$ | 0.205 | 0.130 |
| | Day x Naloxone x Sex | $F_{1,12} < 1$ | | |
| | Day x Morphine x Naloxone | $F_{1,12} < 1$ | | |
| | Day x Morphine x Naloxone x Sex | $F_{1,12} < 1$ | | |
| Figure S3F: Change in event amplitude: after-before naloxone injection | Morphine (between) | $F_{1,12} = 1.28$ | 0.196 | 0.135 |
| | Naloxone (between) | $F_{1,12} = 1.87$ | 0.200 | 0.133 |
| | Morphine x Naloxone | $F_{1,12} = 2.00$ | 0.182 | 0.143 |
| | Sex (between) | $F_{1,12} = 1.15$ | 0.305 | 0.087 |
| | Sex x Morphine | $F_{1,12} = 1.14$ | 0.306 | 0.087 |
| | Sex x Naloxone | $F_{1,12} < 1$ | | |
| | Sex x Morphine x Naloxone | $F_{1,12} < 1$ | | |
| | Day (within) | $F_{1,12} < 1$ | | |
| | Day x Morphine | $F_{1,12} < 1$ | | |
| | Day x Naloxone | $F_{1,12} = 1.45$ | 0.252 | 0.108 |
| | Day x Sex | $F_{1,12} < 1$ | | |
| | Day x Morphine x Sex | $F_{1,12} < 1$ | | |
| | Day x Naloxone x Sex | $F_{1,12} < 1$ | | |
| | Day x Morphine x Naloxone | $F_{1,12} = 1.19$ | 0.296 | 0.090 |
| | Day x Morphine x Naloxone x Sex | $F_{1,12} < 1$ | | |

| Figure | Factor | F-value | p-value | $\eta_p^2$ |
|---|---|---|---|---|
| Figure S4A: Distance travelled during interrupted morphine exposure in *Oprm1* KO mice | Naloxone (between) | $F_{1,10} < 1$ | | |
| | Sex (between) | $F_{1,10} = 3.88$ | 0.077 | 0.279 |
| | Naloxone x Sex | $F_{1,10} = 1.29$ | 0.283 | 0.114 |
| | Day (Within) | $F_{1,10} = 3.29$ | 0.100 | 0.248 |
| | Day x Naloxone | $F_{1,10} < 1$ | | |
| | Day x Sex | $F_{1,10} = 1.73$ | 0.218 | 0.147 |
| | Day x Naloxone x Sex | $F_{1,10} < 1$ | | |
| Figure S4B: Thermal antinociception during interrupted morphine exposure in *Oprm1* KO mice | Naloxone (between) | $F_{1,10} < 1$ | | |
| | Sex (between) | $F_{1,10} < 1$ | | |
| | Naloxone x Sex | $F_{1,10} < 1$ | | |
| | Day (within) | $F_{1,10} = 1.07$ | 0.325 | 0.097 |
| | Day x Naloxone | $F_{1,10} = 1.61$ | 0.233 | 0.139 |
| | Day x Sex | $F_{1,10} < 1$ | | |
| | Day x Naloxone x Sex | $F_{1,10} < 1$ | | |
| Figure S5B: Locomotor activity during continuous iPrecio Infusion | Morphine (between) | $F_{1,24} = 27.71$ | <0.001* | 0.536 |
| | Sex (between) | $F_{1,24} < 1$ | | |
| | Morphine x Sex | $F_{1,24} < 1$ | | |
| | Day (within) | $F_{1,24} = 13.49$ | 0.001* | 0.360 |
| | Day x Morphine | $F_{1,24} = 16.45$ | <0.001* | .407 |
| | Day x Sex | $F_{1,24} < 1$ | | |
| | Day x Morphine x Sex | $F_{1,24} < 1$ | | |
| Figure S5C: Locomotor activity during discontinuous iPrecio Infusion | Morphine (between) | $F_{1,16} = 32.05$ | <0.001* | 0.667 |
| | Sex (between) | $F_{1,16} < 1$ | | |
| | Morphine x Sex | $F_{1,16} < 1$ | | |
| | Day (within) | $F_{2,32} < 1.90$ | 0.166 | 0.106 |
| | Day x Morphine | $F_{2,32} < 1.70$ | 0.199 | 0.096 |
| | Day x Sex | $F_{2,32} < 1$ | | |
| | Day x Morphine x Sex | $F_{2,32} < 1$ | | |
| Figure S5C: Change in locomotor activity (Last – First day) | Group (between) | $F_{2,42} = 17.49$ | <0.001* | 0.454 |
| | Sex (between) | $F_{1,42} < 1$ | | |
| | Group x Sex | $F_{2,42} <1$ | | |
| Figure S6C: Change in Average Fluorescence: After - Before Morphine Challenge | Group (between) | $F_{2,14} = 3.62$ | 0.054 | .341 |
| | Sex (between) | $F_{1,14} < 1$ | | |
| | Group x Sex | $F_{2,14} < 1$ | | |

*Remaining supplementary tables are provided in Excel format.*

**Table S3: Differentially expressed genes (DEGs)** calculated from pair-wise comparisons in the nucleus accumbens and dorsal striatum. Due to limitations in file size for supplemental materials, gene lists are truncated at a nominal p-value of 0.05.

**Table S4: IPA Canonical Pathways.** Lists of all the pathways identified by IPA from the DEGs. Each comparison for each brain region is on separate tabs.

**Table S5: IPA Upstream analysis.** List of all genes identified by IPA as upstream regulators of DEGs. Each comparison for each brain region is on separate tabs.

**Table S6: WGCNA Module Connectivity.** List of all the genes identified in the Magenta, Turquoise and Lime modules and the connectivity for all genes within each module.
